# The king of stress? Exploring the physiological resilience and resistance of adult king penguins to chronic glucocorticoid exposure

**DOI:** 10.64898/2026.07.31.742038

**Authors:** Anais Cotton, Vincent A. Viblanc, Sandra Avril, Lucie Abolivier, Emilie Raymond, Jean-Patrice Robin, Pierre Bize, Pierrick Blanchard, Antoine Stier

## Abstract

To better understand how animals cope with increasingly variable and challenging environments, there is a need to study how prolonged exposure to elevated glucocorticoid hormones (i.e. one mediator of the stress response) affects their physiology. While glucocorticoid elevation is known to increase oxidative stress and accelerate cellular ageing, there is evidence that king penguins (*Aptenodytes patagonicus*) can prevent oxidative stress during acute stress exposure, suggesting that species may differ in their sensitivity to glucocorticoids’ downstream negative effects. As king penguins thrive in a seemingly harsh environment, we hypothesized that they may be able to limit the deleterious effects usually associated with chronic glucocorticoid elevation, either through resistance (i.e. prevention of downstream negative effect) or resilience (i.e. rapid recovery following transient negative effect). To test this hypothesis, we experimentally elevated corticosterone levels in incubating king penguins and quantified treatment effects on a suite of physiological traits at multiple time points across incubation and early chick-rearing, up to ca. 2 months after implantation. Corticosterone-treated individuals showed a prolonged increase in corticosterone and decrease in body condition, confirming our treatment likely mimicked sustained stress exposure. Heterophil-to-lymphocyte ratio was only increased transiently, and there was no clear evidence that treatment influenced oxidative stress or telomere length maintenance. Plasma energy metabolites were mainly affected early after implantation, with rapid recovery over time. Overall, our results suggest that adult king penguins show at least moderate resistance and resilience to chronic corticosterone elevation, especially in preventing cellular integrity loss, though at-sea physiological effects remain to be determined.

## Introduction

In natural environments, animals are constantly facing abiotic and biotic challenges requiring rapid, sometimes sustained, adjustments in their physiology and behaviour to survive and reproduce. In vertebrates, the highly coordinated stress responses are key for responding flexibly and adaptively to unpredictable conditions (McEwen, 1998; Sapolsky et al., 2000; Wingfield & Romero, 2000; Wingfield & Kitaysky, 2002). The vertebrate stress response is primarily mediated by the sympathetic–adrenal–medullary (SAM) and hypothalamic–pituitary–adrenal (HPA) axes. Activation of the SAM axis promotes immediate “fight-or-flight” responses through the rapid release of catecholamines (e.g., adrenaline and noradrenaline), resulting in increased cardiac output, rapid mobilization of glucose, and heightened alertness (Cannon, 1915; Romero & Butler, 2007; Goldstein, 2010). By contrast, activation of the HPA axis leads to the secretion of glucocorticoids (GC) into circulation, which exert slower but longer-lasting effects on energy balance, metabolism, immune function, and other physiological processes that help sustain responses to prolonged or repeated stressors (Sapolsky et al., 2000; Wingfield & Romero, 2000; Romero, 2004; Romero & Butler, 2007). Beyond their role in stress responses, GCs are metabolic hormones that regulate up to 10% of the genome (Le et al., 2005), thereby orchestrating broad physiological reorganisation through energy allocation, immune modulation, and behavioural adjustments. Because they integrate baseline energetic regulation with longer-term stress physiology, glucocorticoids are more commonly used than catecholamines in ecological and evolutionary studies, as SAM-mediated responses are usually too rapid and transient to be reliably quantified in free-living animals. Consequently, GC measurements have become one of the most widely used proxies of stress physiology in wild vertebrate populations (Wingfield et al., 1998; Sapolsky et al., 2000; McEwen & Wingfield, 2003; Romero, 2004; Bonier et al., 2009; Sheriff et al., 2011; MacDougall-Shackleton et al., 2019).

In situations of acute stress, elevated GCs favour the redirection of resources from non-essential functions (e.g. reproduction) toward immediate survival. Enhanced vigilance, increased energy mobilization, and increased responsiveness to external threats all contribute to reducing mortality risk during encounters with predators (Sapolsky et al., 2000; Clinchy et al., 2013; Voellmy et al., 2014). However, chronic elevations of GCs are usually leading to marked physiological costs. At cellular and molecular levels, the deleterious effects of GCs may be mediated by increased oxidative stress, accumulating damage to lipids, proteins, and DNA, impaired repair mechanisms, and accelerated telomere shortening (Haussmann & Marchetto, 2010; Costantini et al., 2011; Monaghan & Haussmann, 2015). At the organismal level, chronically elevated GCs may disrupt energy balance and homeostasis by increasing metabolic demands, leading to prolonged mobilisation of energy reserves and reduced body condition (Sapolsky et al., 2000; Remage-Healey & Romero, 2001; Magomedova & Cummins, 2015). Prolonged exposure to high GC levels may, as a result, compromise growth, reproduction and immune defence, leading to lower fitness (McEwen, 1998; Sapolsky et al., 2000; Romero & Wikelski, 2001; Wingfield & Sapolsky, 2003; Romero, 2004; Romero & Butler, 2007; Breuner et al., 2008; Cyr & Romero, 2009; Angelier et al., 2010; Gouin, 2011). Therefore, chronic GC elevation has often been interpreted as a potential indicator of dysregulated stress responses in wild vertebrates, although this view is debated in ecological contexts where sustained stress responses may also reflect adaptive physiological adjustments (Sapolsky et al., 2000; McEwen, 2001; Romero & Butler, 2007; Sapolsky, 2021; Boonstra, 2013).

However, empirical evidence linking chronic GC elevation to fitness remains equivocal (reviewed in Bonier et al., 2009). Correlative studies based on natural variation in GCs have reported a wide range of relationships with fitness, including strong detrimental costs, weak or absent associations, or even positive links with survival or reproduction (Romero & Wikelski, 2001; Robin et al., 2001; Groscolas et al., 2008; Bonier et al., 2009; Schoenle et al., 2018). In addition, experimental studies manipulating glucocorticoid levels through implants or exogenous administration have also produced heterogeneous results, ranging from negative to neutral or context-dependent effects (Blas et al., 2006; Breuner et al., 2008; Angelier et al., 2009; Breuner et al., 2013). These inconsistencies raise questions about the factors determining the negative, neutral or positive effects of prolonged exposure to elevated GCs.

Two majors, and potentially interacting, factors are thought to shape the outcomes of prolonged exposure to elevated GCs: life-history strategy and environmental conditions. On one hand, life-history theory predicts that long-lived species should prioritize survival over reproduction under stress, because future reproductive opportunities have high fitness value. Consequently, chronic GC elevation may promote resource allocation toward self-maintenance at the expense of reproduction, reflecting life-history trade-offs in stress physiology (Stearns, 1992; Robin et al., 2001; Ricklefs & Wikelski, 2002; Wingfield & Sapolsky, 2003; Groscolas et al., 2008; Angelier & Wingfield, 2013; Sapolsky, 2021). Such reallocation may enhance short-term maintenance or immediate survival but can incur longer-term physiological and fitness costs if sustained over time. Consistent with this temporal framework, a phylogenetically controlled meta-analysis showed that elevated GCs are associated with reduced survival in long-lived species, suggesting that chronic elevation may be particularly costly over extended timescales, whereas short-lived species appear more tolerant to sustained GC exposure (Schoenle et al., 2021). On the other hand, environmental conditions further modulate these outcomes by influencing both the magnitude and consequences of HPA axis activation. The intensity, frequency, and predictability of stressors shape physiological responses such that elevated GCs may enhance survival and fitness in harsh or high-risk environments but become detrimental in less challenging environments (Cabezas et al., 2007; Patterson et al., 2014; Jaatinen et al., 2014; Jimeno et al., 2018; reviewed in Schoenle et al., 2018; Zimmer et al., 2020). For example, a three-year study of blue tits (*Cyanistes caeruleus*) found that parental baseline corticosterone levels were elevated only in the year when environmental conditions were harsher. In that year, parents with higher baseline corticosterone also fledged more offspring, whereas such a relationship was not present in more favourable years (Henderson et al., 2017). These context-dependent effects suggest that the fitness consequences of GC elevation are not fixed but vary with both life-history strategy, environmental conditions and their interaction. This variability implies that species may buffer the potential costs of chronic GC elevation through resistance, i.e., the ability to maintain physiological and behavioural functioning despite prolonged glucocorticoid elevation, or through resilience, i.e. the capacity to rapidly restore normal physiological states following GC-induced perturbations (Wingfield & Sapolsky, 2003; Vitousek et al., 2019; Lane et al., 2025). Both strategies may allow individuals to cope with the potentially deleterious effects of chronic stress and are likely to be particularly important for species living in harsh environments, where frequent or prolonged stressors demand sustained stress responsiveness while minimising fitness costs (Wingfield & Sapolsky, 2003). Investigating resistance and resilience requires experimental approaches that quantify not only the magnitude but also the temporal dynamics of physiological responses to chronically elevated GC exposure. Despite growing interest in these coping mechanisms, such experimental tests remain rare, especially in the wild (but see for instance O’Connor et al., 2009; Murone et al., 2016; Names et al., 2021).

King penguins (*Aptenodytes patagonicus*) provide an interesting model to address this question. These long-lived seabirds breed on sub-Antarctic islands and experience repeated and prolonged challenges during reproduction including: extended periods of fasting (Groscolas et al., 2001), intense social aggressive interactions at breeding sites (Côté, 2000), demanding foraging conditions at sea (Brisson-Curadeau et al., 2024; Lemonnier et al., 2025), and an extended reproductive cycle on land (Weimerskirch et al., 1992). Therefore, we hypothesise that king penguins may have evolved specific physiological mechanisms that confer resistance and/or resilience to prolonged GC elevation, enabling them to thrive in harsh conditions while minimizing negative effects. Supporting this idea, previous work has shown that adult king penguins avoid the typical oxidative cost of acute stress exposure and associated GC increase (Majer et al., 2019) through upregulation of antioxidant defences (Stier et al., 2019).

Here, we experimentally tested whether adult king penguins exhibit resistance and/or resilience to a prolonged elevation in GC levels. We elevated circulating plasma corticosterone (CORT), the primary GC in birds, in free-living incubating males and females using subcutaneous CORT-releasing pellets that provided sustained hormone release. We analysed multiple physiological systems expected to reflect GC-induced costs. Chronic stress markers were assessed through plasma CORT levels and Heterophil/Lymphocyte (H/L) ratio, two widely used indicators of prolonged activation of the HPA axis in wild vertebrates (Romero, 2004; Davis et al., 2008; Sheriff et al., 2011). Oxidative stress was measured using markers of oxidative damage and antioxidant defences (Costantini et al., 2011; Haussmann & Marchetto, 2010; Monaghan & Haussmann, 2015). Relative telomere length, a frequently used biomarker of cumulative physiological stress and future survival prospects, was used as a proxy of cellular ageing (Haussmann & Marchetto, 2010; Monaghan & Haussmann, 2015). Energy metabolism was characterised by circulating metabolites and heart rate, both of which are commonly used to quantify GC-induced changes in energy allocation and autonomic function (Sapolsky et al., 2000; McEwen & Wingfield, 2003; Cyr & Romero, 2009).

If king penguins show resistance to prolonged GC elevation, we predict little evidence of physiological costs. In this case, elevated GC levels should not result in oxidative damage as a compensatory increase in antioxidant defences might be expected to regulate any increase in reactive oxygen production resulting from chronically elevated GC (Manoli et al., 2007). Under this scenario, body condition and proxies of energy metabolism would not be affected either. If king penguins show resilience to prolonged GC elevation, we would expect oxidative stress, metabolic changes, and body condition to be transiently affected, followed by a rapid recovery to baseline levels despite continued exposure. In the context of both resistance and resilience, we would not expect deleterious effects on telomere lengths as they are a marker of cumulative stress exposure and future fitness prospects (Wilbourn et al., 2018; Eastwood et al., 2019). Together, these predictions allow us to test whether king penguins show resistance and/or resilience to chronic GC elevation by examining both the magnitude and the temporal dynamics of physiological responses to chronically elevated circulating GCs.

## Methods

### Study site and species

This study was conducted during the 2018–2019 breeding season in a king penguin colony of approximately 22,000 breeding pairs located at La Baie du Marin on Possession Island, Crozet Archipelago, southern Indian Ocean (Barbraud et al., 2020). At the beginning of the breeding season, partners alternate between parental duties (incubation and early chick guarding) on land and foraging trips at sea (Weimerskirch et al., 1992). Males typically undertake the first incubation shift while their partner forages at sea. After about 15 days, females return to relieve their partner, and the pair alternates shifts for a total incubation period of approximately 53 days, and a chick-guarding phase of approximately one month (Weimerskirch et al., 1992).

### Experimental procedure

We monitored 49 breeding pairs from the onset of courtship in early November through the end of the chick-guarding phase, approximately 2.5 months later. The 49 breeding pairs were randomly assigned to two experimental groups: female-treated pairs (n = 23) and male-treated pairs (n = 26). For each pair, we treated only one partner (male or female) at a time. Each adult was individually identified using hair dye (Lemonnier et al., 2025) marking on the breast feathers. Both experimental groups were sampled over the same period, ensuring that individuals experienced comparable environmental conditions both on land and at sea throughout the study. In female-treated pairs: females received either a CORT (NG-111, 50 mg, 90-day release; n = 12) or a placebo (control, C) implant (NC-111, identical composition without hormone; n = 11) (8 mm × 3 mm; Innovative Research of America, Sarasota, FL, USA). Implants were inserted subcutaneously three days after females had returned from their first foraging trip at sea, i.e. first incubation shift (Fig. 1). In male-treated pairs: 13 males received a CORT implant and 13 received a C implant. Conditions were identical to those of females, except that males were implanted during their second incubation shift to match the fasting duration of treated females (approximately 3 days), ensuring a comparable physiological state between sexes at implantation, as males had already been fasting for approximately two weeks prior to the onset of their first incubation shift (Weimerskirch et al., 1992).

**Fig. 1.**
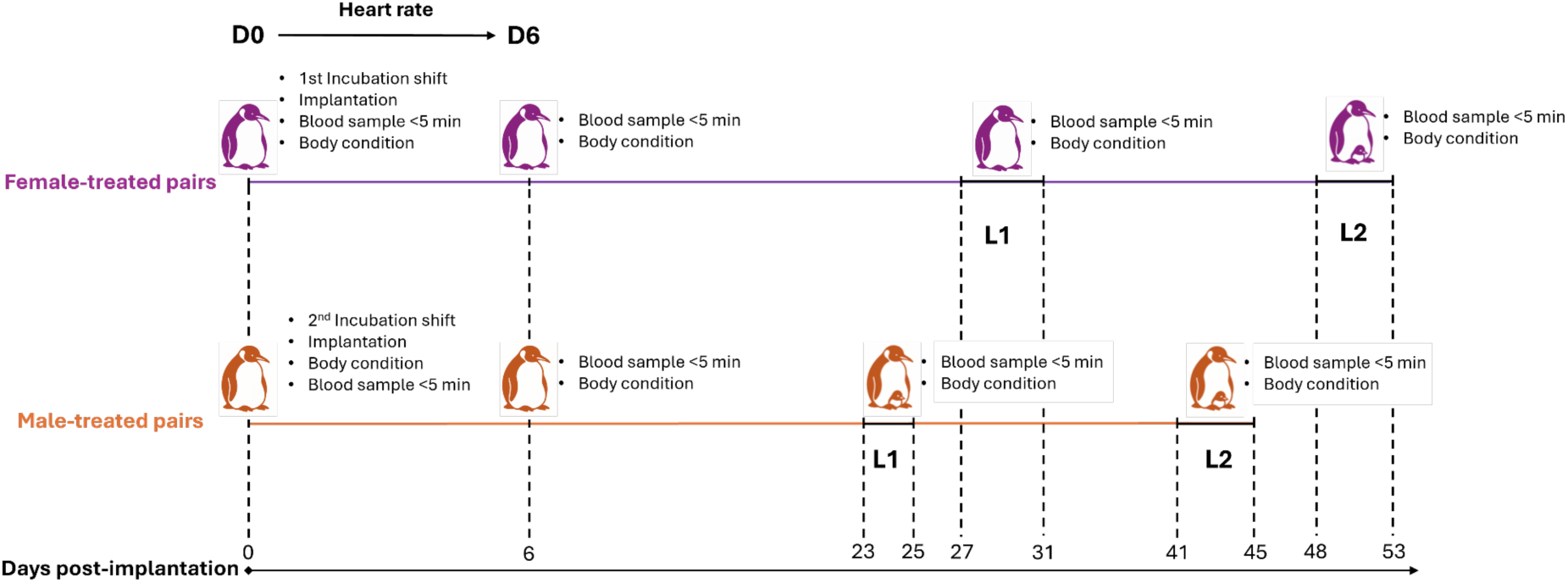
Schematic representation of the study protocol and timing of captures following corticosterone or placebo pellet implantation in breeding king penguin adults. Individuals were captured at implantation (Day 0, D0), six days after implantation (D6), after the first post-implantation period at sea and return on land (L1), and after the second post-implantation period at sea and return on land (L2). Blood samples and thorax circumference measurements were collected at each capture event, whereas complete morphological measurements were obtained at D0 only. The figure also shows the 95% confidence intervals for the number of days elapsed since implantation when individuals returned on land after the first (L1) and second (L2) post-implantation foraging periods (solid black lines). Brooded chicks are represented schematically to indicate when the chick-rearing phase started on average.

On implantation day (D0), adults were captured by hand while incubating and fitted with a hood to reduce stress. A blood sample was collected within 5 minutes to measure pre-treatment physiological parameters including baseline CORT levels (see Physiological assays). Under local anaesthesia (ca. 0.5 mg/kg xylocaine and 0.0001 mg/kg adrenaline; Aspen Pharma), a veterinarian (SA) inserted the implant subcutaneously into the upper back. The ∼15 mm incision was closed with three sterile staples (SurgiClose™), and a prophylactic antibiotic (cephalexin, ∼50 mg/kg; Rilexine®, Virbac) was administered. Six days post-implantation (D6), all treated individuals were recaptured, staples removed, and no signs of infection or implant rejection were observed. A blood sample was collected following the same protocol as at D0. Treated individuals were recaptured twice for measurements and blood sampling (<5 min) after their first (L1) and second (L2) foraging trips at sea and return on land. Sampling always occurred after 3 days of fasting, on the third day after returning to land. For females, L1 occurred 22–40 days post-implantation, and L2, 41–60 days post-implantation. For males, L1 occurred 18–28 days post-implantation, and L2, 35–55 days post-implantation (Fig. 1).

### Physiological assays

#### 1. Stress markers

##### Corticosterone (CORT)

Baseline corticosterone (CORT) concentrations were measured from 25 μL of plasma using a commercial enzyme immunoassay kit (Corticosterone Enzyme Immunoassay Kit, Arbor Assay, USA), following the manufacturer’s instructions and as previously applied in king penguins (Lewden et al., 2024). Intra-plate coefficient of variation (CV) was 7.3 ± 0.8% (mean ± SE), and inter-plate CV was 13.9%. Technical repeatability based on duplicates was high (*R* = 0.98, p < 0.001).

##### Heterophil/Lymphocyte ratio (H/L)

The heterophil-to-lymphocyte ratio (H/L) is widely used as an indicator of chronic stress and immune function in birds, including king penguins (Lemonnier et al., 2026), as prolonged glucocorticoid exposure induces an increase in circulating heterophils and a decrease in lymphocytes (Gross & Siegel, 1983; Davis et al., 2008). Blood smears were prepared in the field on freshly collected blood, air-dried for 24–48 h, and stained using a RAL555 kit (RAL Diagnostics, Martillac, France). Differential white blood cell counts were performed by the same experimenter (SA). Technical precision based on 10 repeated samples was 17.3 ± 2.8%.

#### 2. Oxidative stress and telomere length

We quantified oxidative stress by combining markers of oxidative damage and antioxidant defences and assessed relative telomere length as a biomarker of cellular ageing (Blackburn, 2000; Monaghan, 2010).

##### Oxidative damage

Oxidative damage to DNA was evaluated using 8-hydroxy-2’-deoxyguanosine (8-OHdG), measured with a DNA damage ELISA kit (Arbor Assay, USA). Plasma 8-OHdG (dilution 1/20) reflects whole-body 8-OHdG excretion, which integrates both DNA damage and repair processes (Stier et al., 2014b; Stier et al., 2019). Values are expressed as ng/mL (intra-plate CV = 5.8 ± 0.9%, inter-plate CV = 4.1%, R = 0.95, p < 0.001). DNA 8-OHdG was measured on genomic DNA extracted from blood cells, reflecting DNA damage incorporated into the nuclear DNA of blood cells (Stier et al., 2019). One microgram of DNA was enzymatically digested as described in Stier et al. (2014a) prior to ELISA (expressed as pg/μg DNA; intra-plate CV = 9.5 ± 1.8%, inter-plate CV = 6.5%, R = 0.80, p < 0.001).

Reactive oxygen metabolites (ROMs) in plasma were measured using the d-ROM test (5 µL of plasma, Diacron International, Italy), following manufacturer instructions and previous work in king penguins (Stier et al., 2014b; Viblanc et al., 2016; Stier et al., 2019). This assay primarily quantifies hydroperoxides (ROOH) as a marker of potential oxidative damage and has been widely used in birds (Costantini, 2016). ROMs are expressed as mg H₂O₂ equivalent/dL (intra-plate CV = 14.3 ± 1.6%, inter-plate CV = 12.6%, R = 0.93, p < 0.001).

##### Antioxidant defences

We measured both enzymatic and non-enzymatic antioxidant defences. Superoxide dismutase (SOD) activity was quantified in red blood cell lysate (dilution ca. 1:200) using the RANSOD kit (Randox Laboratory, UK), following manufacturer instructions and previous work in king penguins (Stier et al., 2019). SOD values were corrected for lysate actual concentration by including BCA-derived protein quantification as covariate in the statistical models. SOD activity is expressed as U·mL⁻¹ (intra-plate CV = 17.0 ± 3.4%, inter-plate CV = 15.6%, R = 0.78, p < 0.001). Glutathione peroxidase (GPx) activity in red blood cells (dilution ca. 1:50) was measured using the RANSEL assay kit (Randox Laboratory, UK), as in Stier et al. (2019). GPx values were similarly corrected, with BCA protein measurements. GPx activity is expressed as U·L⁻¹ (intra-plate CV = 10.3 ± 1.2%, inter-plate CV = 22.5%, R = 0.87, p < 0.001).

The non-enzymatic antioxidant capacity (OXY) of the plasma (dilution 1:100) was measured using the OXY-Adsorbent test (Diacron International, Italy), following manufacturer instructions and previous applications in king penguins (Stier et al., 2014b). This assay quantifies the ability of non-enzymatic antioxidants (e.g. vitamins, carotenoids, flavonoids, thiols) to neutralize a massive oxidative challenge induced by hypochlorous acid (HClO). OXY is expressed as μM HClO neutralised·L⁻¹ (intra-plate CV = 7.7 ± 1.0%, inter-plate CV = 9.0%, R = 0.81, p < 0.001).

##### Relative telomere length (rTL)

Genomic DNA was extracted from 4 μL of packed blood cells using the NucleoSpin® Blood QuickPure kit (Macherey-Nagel), following manufacturer instructions. DNA concentration and purity (260/280 and 260/230 ratios) were checked; samples not meeting quality criteria (concentration > 50 ng/μL; ratios > 1.8) were re-extracted when possible or excluded. DNA integrity was confirmed in a subset of samples by gel electrophoresis. DNA was stored at −80°C until qPCR and diluted to 0.6 ng/μL just before assays. Relative telomere length was measured by qPCR as previously done in this species (Stier et al., 2014b, 2019), with minor modifications. Telomere (Tel1b/Tel2b) and single-copy nuclear gene (RAG1) reactions were run on separate plates using Sensifast SYBR® No-ROX Mix (Bioline). RAG1 was chosen as a reference gene (primers verified on *Aptenodytes forsteri* genome). Each sample was run in duplicate, and samples from the same individual were placed on the same plate. A pooled DNA sample was used as a reference (ratio = 1) on every plate, and one inter-plate standard sample was also included. qPCR was performed on a Mic qPCR instrument (Bio Molecular Systems) with a two-step cycling protocol (95°C for 3 min; then 25 or 38 cycles of 95°C for 5 s and 60°C for 25 s, with fluorescence read at 60°C for telomere and RAG1 reactions, respectively). Amplification efficiencies were estimated using both standard curves and the LinReg algorithm (RAG1: 97.4 ± 0.9%; Tel: 97.7 ± 0.2%). rTL was calculated as (1 + Ef_Tel)^ΔCqTel / (1 + Ef_RAG1)^ΔCqRAG1, where Ef is amplification efficiency and ΔCq the difference between the Cq of the reference and the sample. Intra-plate CV for rTL was 11.4 ± 1.5%, inter-plate CV 13.1%; repeatability within plates (based on duplicates) was R = 0.84, p < 0.001, and inter-plate repeatability based on two repeated plates was R = 0.91, p < 0.001. Adjusted within-individual repeatability of rTL was R = 0.66 (95% CI: [0.51; 0.78], p < 0.001), which is high relative to typical qPCR-based telomere studies (Kärkkäinen et al., 2021).

#### 3. Body condition, heart rate and plasma energy metabolites

##### Body condition

At implantation (D0), thorax circumference (a validated proxy for body mass in this species; Viblanc et al., 2012a), flipper length (mean of left and right), and beak length were measured to the nearest 1 mm. Thorax circumference was then measured at each subsequent capture event. Structural size was quantified using a principal component analysis (PCA) performed on flipper and beak length following established protocol (Saraux et al., 2011). The first principal component (PC1) explained 73.2 % of the variance and was used as a structural size index (SSI), defined as PC1 = 0.25 × beak length + 0.75 × flipper length. Thorax circumference was regressed on PC1 (*F*_1,177_ = 13.55, p < 0.001), and residuals were used as an index of body condition. Body condition could not be calculated for one individual because size was never measured for this bird.

##### Heart rate measurements

Heart rate (HR) was recorded using external HR loggers (Polar® RS800, Polar Electro Oy, Kempele, Finland), following established protocols in king penguins (Groscolas et al., 2010; Viblanc et al., 2012b; Lemonnier et al., 2024). At implantation, the transmitter (weighing <1% of body mass) was attached to the bird’s back with Tesa® tape, and the receiver was positioned on a plastic flipper band. HR was recorded every 5 s for three consecutive 24 h periods during each recording session. Each individual underwent two recording sessions, yielding up to six consecutive 24 h periods of post-implantation heart rate (HR) data, with a battery change required between sessions. HR data were analysed using Polar Pro Trainer 5 software and expressed in beats per minute (BPM). To characterise energy metabolism, we calculated: (i) mean HR as the average of all HR measurements over 24 h and (ii) resting HR as the lowest 10-minute mean HR within the same day. On D0, HR was elevated immediately after handling, and the first hour of data was excluded (Viblanc et al., 2012b). On D4, logger replacement induced handling-related increases in HR and data loss; this day was therefore excluded from analyses. Data of insufficient quality (e.g. due to signal loss or very high noise) were removed from the analysis.

##### Plasma energy metabolites

Plasma metabolites were measured as indicators of energy availability and metabolic adjustments under chronic CORT elevation. Glucose, triglycerides, uric acid, and lactate were analysed using Randox assay kits (Randox Laboratory, UK). Glucose is the main circulating carbohydrate, reflecting immediate energy availability and is known to be modulated by GCs during stress to support energy-demanding processes (Bernard et al., 2003; Braun & Sweazea, 2008). Triglycerides levels are an index of lipid reserve mobilisation, particularly relevant during prolonged fasting or sustained stress (Sapolsky et al., 2000). Uric acid levels reflect protein catabolism associated with amino acid breakdown, which can be enhanced under chronic stress (Sapolsky et al., 2000). Lactate levels indicate a shift towards anaerobic metabolism, typically associated with acute stress and increased oxygen demand (Le Maho et al., 1992). Technical precision and repeatability were as follows: Glucose: intra-plate CV = 10.7 ± 0.6%, inter-plate CV = 8.2%; R = 0.66, p < 0.001; Triglycerides: intra-plate CV = 7.3 ± 0.6%, inter-plate CV = 16.9%; R = 0.89, p < 0.001; Uric acid: intra-plate CV = 8.4 ± 0.6%, inter-plate CV = 11.7%; R = 0.91, p < 0.001; Lactate: intra-plate CV = 7.1 ± 0.5%, inter-plate CV = 14.5%; R = 0.96, p < 0.001.

### Statistical analyses

We performed all statistical analyses in R, version 4.4.2. We divided the data into two databases: pre-implantation measurements (D0) and post-implantation measurements (D6, L1, and L2). Assumptions for linear and generalised linear mixed models (LMMs) (overdispersion, homoscedasticity, independence and normality of models’ residuals) were checked with packages ‘car’ (Fox & Weisberg, 2019) and ‘DHARMa’ (Hartig, 2022). Model assumptions were checked by visual inspection of residuals and confirmed using statistical tests (p > 0.05). As LMMs are generally robust to moderate violations of these assumptions (Schielzeth et al., 2020), any significant deviation is explicitly reported in the results. Results of type III analysis of variance from mixed models have been obtained through the anova() function and the package lmerTest (Kuznetsova & Christensen, 2017), while model estimates are provided in Tables S1-S5. Sample size varies slightly between parameters because of missing data or failed laboratory assays and are fully reported in ESM Tables S1-S5.

### Pre-implantation measurements (D0)

As a first step, we tested whether physiological parameters at pre-implantation differed between treated and control birds using linear models (LM) and found no significant differences among them (all *F*_1,47_ < 3.45, p > 0.070). Additionally, we tested whether pre-implantation physiological values were related to blood sampling duration (mean = 2.44 min, 95% CI: 2.11-2.67) using a LM with sampling duration as a fixed effect. Only lactate levels increased significantly with sampling duration (*F*_1,43_ = 15.102, p < 0.001), consistent with Viblanc et al. (2018), reporting a rise after 3 minutes of capture. Since we use pre-implantation levels as a covariate to control for initial individual differences when assessing the treatment effect, pre-implantation lactate levels were corrected for blood sampling duration (i.e. residuals from a linear regression). Other physiological parameters showed no significant relationship to sampling duration (*F*_1,44_ ≤ 3.643, p ≥ 0.063). CORT, H/L ratio, dROMs, UA were log-transformed to meet model assumptions.

### Post-implantation measurements (D6, L1, L2)

We investigated whether physiological parameters and individual body condition at post-implantation differed between control and treated birds, using linear mixed effect regression models (LMM) with individual identity and assay plate or experimenter identity specified as random factors. We initially included the period (D6, L1, L2), sex, treatment, and all two-way interactions as fixed factors. To account for the influence of initial physiological values at pre-implantation on post-implantation parameters, we included each parameter’s pre-implantation value as a covariate in the models, except for 8-OHdG DNA damage and 8-OHdG plasma levels, as only a small number of individuals had pre-implantation data for these markers (n = 20 and n = 17 respectively). Similarly, we included blood sampling duration (mean = 2.39 min, 95% CI: 2.17-2.61) as a covariate to avoid potential confounding effects on physiological parameters, as longer blood sampling durations (even when < 5 minutes) can lead to changes in the measured parameters (Viblanc et al., 2018). We tested the two-way interactions between treatment, period and sex (Treatment × Period, Treatment × Sex, and Sex × Period) to evaluate whether treatment effects varied over time or were modulated by sex. In particular, we focused on Treatment × Period as it captures temporal dynamics of the treatment response, and on Treatment × Sex as it allows testing for sex-specific sensitivity to the treatment, while Sex × Period accounts for potential baseline temporal variation between sexes. We then dropped interaction terms with p > 0.10 (starting with higher-order interactions) in a backward stepwise procedure. We subsequently performed multiple post hoc comparisons tests using Tukey correction using the emmeans R package (Lenth et al., 2019). All physiological parameters, except GPx and OXY, were log-transformed to meet model assumptions.

### Heart rate analysis (D0-D6)

To analyse whether mean HR and resting HR differed between control and treated birds, we used a LMM with individual identity as a random factor. We included treatment group, number of days since implantation (Day, continuous), and sex as fixed factors, and tested the corresponding two-way interactions (Treatment × Day, Treatment × Sex, Sex × Day) to account for potential temporal dynamics of treatment effects, sex-dependent physiological sensitivity, and sex-specific temporal trajectories in stress and metabolic regulation. Only significant interactions were retained in the final models. We initially tested the effect of HR measurement duration as a covariate to account for potential missing data from the full 24-hour record. As this effect was not significant (*F*_1,44_ ≤ 2.823, p ≥ 0.095), it was removed from the final models.

### Effect size calculation

Effect sizes (approximated as Cohen’s *d*) and the associated 95% confidence intervals were calculated from model-derived contrasts using estimated marginal means and were used for visualisation. These were obtained with the eff_size() function from the R package “emmeans”, using the residual standard deviation and associated degrees of freedom from each model.

## Results

### 1. Stress markers

Baseline corticosterone levels of CORT-treated individuals increased significantly in response to the implant. Specifically, CORT levels were 2.58 times higher on D6, 1.57 times higher on L1, and returned close to Placebo bird levels on L2 (Treatment × Period: *F*_2,73.3_ = 16.58, p < 0.001, Table S1A, Fig. 2). There was a transient increase in H/L ratio in CORT-treated birds, showing a 2.14-fold increase on D6 compared to Placebo birds, while values at L1 and L2 returned close to Placebo bird levels (Treatment × Period: *F*_2,122_ = 6.42, p = 0.002, Table S1B, Fig. 2).

**Fig. 2.**
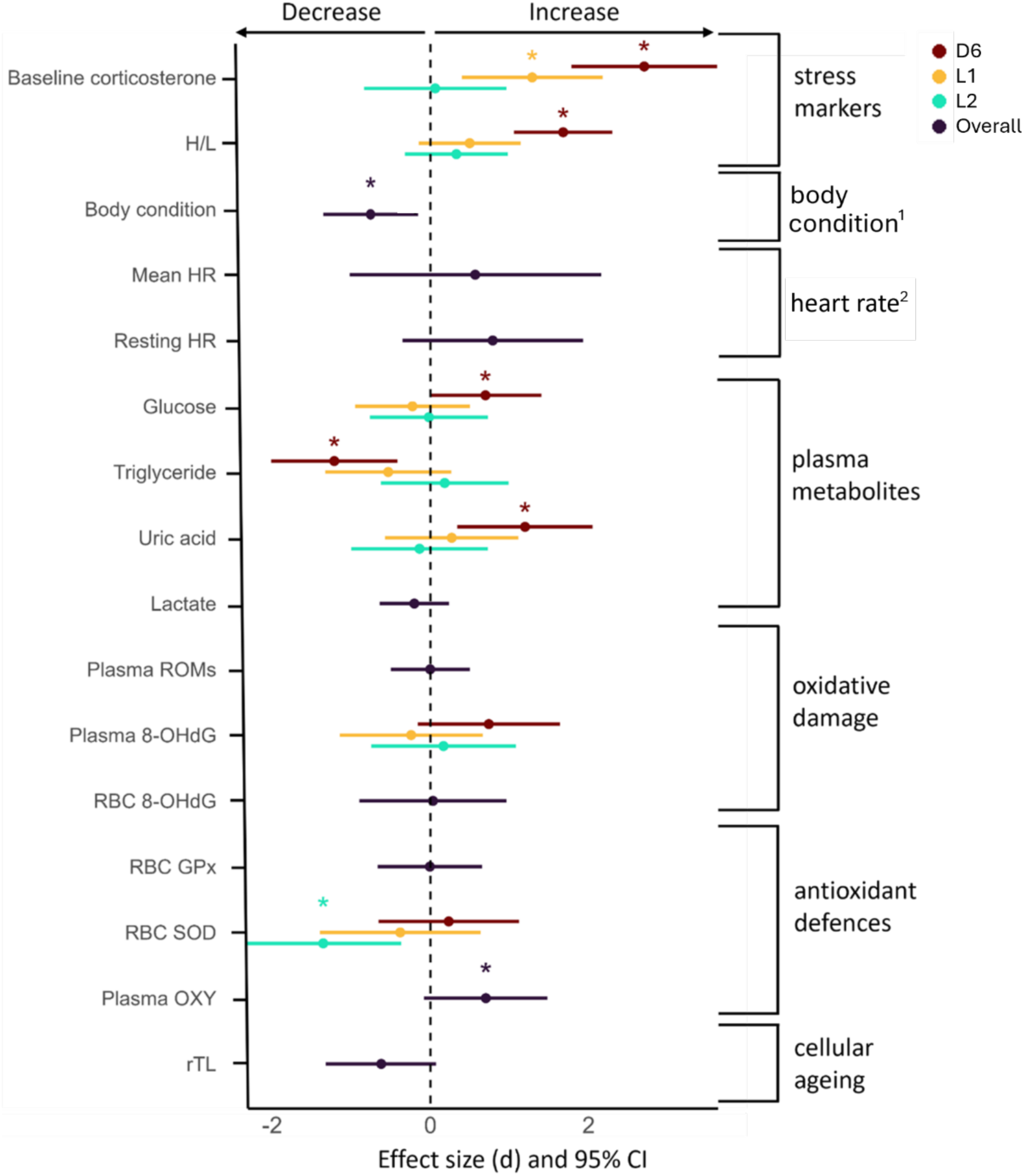
Effects of corticosterone implant on king penguin physiology. Standardized effect sizes (*d*) and 95% confidence intervals of the effect of corticosterone implants are reported (i.e. compared to placebo implants) along with their 95% confidence intervals are shown. When there was some evidence of an interaction between the corticosterone treatment and the time post-implantation (Treatment × Period: p < 0.10), colours indicate the different periods: brown for day 6 after implantation (D6), orange for the first period on land after returning from foraging at sea post-implantation (L1: 18–40 days), and green for the second period on land after returning from the second foraging trip at sea (L2: 35–60 days). If there was no evidence of an interaction between Treatment × Period (p > 0.10), points are shown in deep blue to reflect the main effect of Treatment only. Statistically significant differences (i.e. p < 0.050, see Tables S1-S5 in ESM) between CORT and Placebo according to post-hoc comparisons are marked with a * Superscript ¹ refers to Fig S1 presenting the response for each period, and ² refers to Figures S2, which presents the interaction between Treatment and time over the first six-days post-implantation. Details on the statistical tests and sample sizes for each parameter are provided in Tables S1-S5.

### 2. Body condition and HR

#### Body condition

Individual body condition was lower in CORT-treated birds (Treatment: *F*_1,42.69_ = 6.15, p = 0.017, Table S2, Fig. 2). Yet, although there was no significant Treatment × Period interaction (p = 0.22), the gap between Placebo and CORT birds appeared to widen over time, suggesting a potential tendency for the Treatment effect to increase across the study period (see Fig. S1).

#### Heart rate

Mean HR significantly decreased over the 6 days after implantation (Day: *F*_1,158.63_ = 10.91, p = 0.001, Fig. S2, Table S3A). While mean HR did not differ significantly between Treatment groups (Treatment: *F*_1,42.45_ = 0.74, p = 0.46; Fig. S2, Table S3A), Resting HR decreased more strongly over time in CORT-treated pairs when averaged across sex (β = −2.85 ± 0.59 bpm day⁻¹; approximately −3.4% per day relative to model-estimated baseline resting HR) compared to placebo pairs (β = −0.91 ± 0.61 bpm day⁻¹; approximately −1.0% per day) (Day × Treatment: *F*_1,158.43_ = 5.23, p = 0.024, Fig. S2, Table S3B).

### 3. Plasma metabolites

Linear mixed models revealed significant Treatment × Period interactions for plasma triglycerides (*F*_2,66.47_ = 4.24, p = 0.018) and uric acid levels (*F*_2,71.38_ = 4.19, p = 0.019), and a marginal interaction for plasma glucose (*F*_2,76.41_ = 2.45, p = 0.093). Post-hoc comparisons indicated significant treatment effects at Day 6 on glucose (p = 0.047), triglycerides (p = 0.002) and uric acid (p = 0.002) at Day 6. CORT-treated individuals exhibited a 1.10-fold increase in plasma glucose, a 1.41-fold decrease in plasma triglycerides, and a 1.42-fold increase in uric acid relative to the placebo group (see Fig. 2, Table S4A–C). These effects were short-lived, as metabolite levels did not differ from placebo at L1 and L2 (all p > 0.17). No differences were observed for plasma lactate between treatment groups (*F*_1,104.95_ = 1.11, p = 0.29; Table S4D).

### 4. Oxidative stress & relative telomere length

No markers of oxidative damage differed significantly between CORT-treated and Placebo individuals, neither as a main effect nor in interaction with Period (all p > 0.095, see Table S5A-C, Fig. 2) or Sex (all p > 0.23, see Table S5A-C, Fig. 2). GPx activity did not significantly differ between treatment groups (Treatment: *F*_1,37.63_ = 0.001, p = 0.98, Table S5D). In contrast, SOD activity in RBC was lower in CORT-treated individuals), but only a long-time after implantation at L2 (Treatment × Period: *F*_2,42.59_ = 3.75, p = 0.032; Table S5E, Fig. 2). Plasma non-enzymatic antioxidant capacity was significantly higher in CORT-treated individuals (Treatment: *F*_1,37.34_ = 5.67, p = 0.022, Table S5F). Relative telomere length (rTL) did not differ significantly between treatment groups (Treatment: *F*_1,43.44_ = 3.59, p = 0.065, Table S5G), although CORT-treated individuals showed a 1.09-fold lower rTL compared to placebo (see Fig. 2).

## Discussion

By experimentally increasing circulating CORT levels in free-living king penguins, we tested whether this long-lived seabird exhibited resistance and/or resilience to chronic GC elevation. Our experiment successfully increased bird GC levels over a 3-5 week-period and was associated with a decline in body condition over a 6-8 week-period. Despite this, the downstream effects of increased GCs on heterophil/lymphocyte ratio, oxidative stress, telomere length, plasma metabolites and whole-body metabolism were mostly absent or transient, suggesting a combination of both partial resistance and resilience to chronic GC elevation.

The increase in CORT levels in our study was shorter (ca. 1 month) than expected from using 90-days releasing CORT pellets. Yet, it has already been described that CORT implants usually raise plasma levels on a shorter timeframe than expected in birds, which has been suggested to result from the HPA negative feedback loop reducing endogenous corticosterone production (Dallman et al., 1992, but see Torres-Medina et al., 2018). Despite a shorter CORT elevation period, body condition decreased in the treated group over a 6–8-week period, with a non-significant trend for this effect to increase with time. CORT negative effects on body condition have been well documented across a variety of bird species (e.g. Kitaysky et al., 2007; Angelier et al., 2009; Spée et al., 2011), and these effects can be attributed to two potential processes. The first is an increase in energy expenditure due to GC-induced metabolic activation, which can include elevated basal metabolism, thermogenesis, and locomotor activity (Belthoff & Dufty, 1998; Romero & Wikelski, 2002; Lynn et al., 2003; Landys et al., 2006). Our results on heart rate (i.e. proxy of metabolism) do not support this hypothesis, since CORT-treated individuals showed similar average heart rate than placebos, and even a steeper decline in resting heart rate along the fasting duration on land. The second process leading to a reduction in body condition would be a decrease in energy acquisition. Experiments have shown that elevated CORT levels can affect foraging behaviour and food intake: in Adélie penguins, Spée et al. (2011) found that incubating males spent less time at sea searching for food after CORT implantation and showed a decrease in body condition. In other seabirds, elevated GCs enhance foraging effort but do not necessarily translate into greater energy intake (Kitaysky et al., 2003; Angelier et al., 2007; Angelier et al., 2008). In king penguins, individuals abandoning reproduction during phase 3 of fasting (i.e. a late fasting stage characterized by severe depletion of energy reserves and a shift towards increased protein catabolism) exhibit elevated glucocorticoid levels and show an increased mass gain during the subsequent foraging trip at sea compared to birds still engaged in reproduction (Cherel et al., 1988; Robin et al., 2001). This is probably explained by an increase in energy intake, which is likely the opposite to what happens for our experimental increase in CORT considering the decrease in body condition we observed. Contrary to the pattern observed on body condition, several physiological pathways showed a short-term alteration linked to the CORT treatment followed by a quick return to baseline levels. This pattern is consistent with the concept of stress resilience, whereby efficient regulation of the HPA axis facilitates recovery from GC-induced physiological perturbations (Taff et al., 2018; Zimmer et al., 2019; Vitousek et al., 2019). First, the H/L ratio, a biomarker of chronic stress and immune function, was markedly elevated a few days following implantation, but no longer after the first foraging trip at sea. GCs increase heterophils while decreasing lymphocytes, thereby increasing the H/L ratio. Multiple studies have found a positive association between baseline CORT and the H/L ratio. While the H/L ratio is not directly regulated by the HPA system, its variation usually reflects downstream effects of changes in glucocorticoid levels (Gross & Siegel, 1983; Dhabhar, 2002; Davis et al., 2008). Therefore, the fact that baseline CORT remained elevated after the first foraging trip, while H/L did not, suggests potential resilience mechanisms being able to restore white blood cell proportions despite a chronic elevation in GCs. In support of this, in other bird species, an experimental and sustained elevation of GCs via injections or implants has been associated with a sustained increase in the H/L ratio over a two-week period compared to controls (Mehaisen et al., 2017; Oluwagbenga et al., 2023).

Plasma energy metabolites exhibited a pattern similar to H/L ratio, with a short-term increase in glucose and uric acid, and concurrent decrease in triglycerides, followed by a full recovery after the first foraging trip at sea. CORT affects how birds process carbohydrates, proteins, and lipids, boosting glucose production, mobilising amino acids, and mobilising lipids to meet energy needs (Sapolsky et al., 2000; Remage-Healey & Romero, 2001). In other birds, studies have shown that glucose and uric acid levels tend to spike early during chronic corticosterone treatment, before returning to baseline, consistent with a physiological negative feedback response (Romero et al., 2005; Bize et al., 2010; Ouyang et al., 2013). King penguins, in this context, appear neither more nor less resilient than these other short-lived species (Romero et al., 2005; Bize et al., 2010; Ouyang et al., 2013). In contrast to species where prolonged exposure to GCs generally leads to an increase in circulating triglycerides (Warne et al., 2009; Campbell et al., 2011), king penguins exhibit a transient decrease in triglyceride levels. Similar responses have been observed in other bird species experiencing acute stress (Remage-Healey & Romero, 2001; Butler et al., 2020). This suggests that triglycerides are rapidly converted into free fatty acids (FFA) and used by cells to produce energy, in line with previous findings showing a rapid mobilisation of circulating lipids following acute stress in king penguins (Viblanc et al., 2018), allowing king penguins to meet immediate metabolic demands, a process not sustained over time, suggesting that king penguins are likely resilient in maintaining a balanced lipid metabolism, efficiently mobilising energy when needed without disrupting lipid metabolism over the long-term.

One commonly reported negative effect of elevated CORT is an increase in oxidative stress (Finkel & Holbrook, 2000; Manoli et al., 2007; Monaghan et al., 2009; Costantini et al., 2011; Metcalfe & Monaghan, 2013). For instance, in kestrels (*Falco tinnunculus*), chronic CORT exposure led to a 32% increase in reactive oxygen metabolites compared to controls (Costantini et al., 2008). In broiler chicken (*Gallus gallus domesticus*), a 3-day administration of CORT was enough to induce oxidative stress (Lin et al., 2004). In contrast, our study shows no major changes in oxidative damage or antioxidant defences in response to chronic CORT treatment. This is consistent with previous findings in king penguins showing no increase in oxidative damage in response to acute restraint stress, indicating an apparent resistance to oxidative stress (Stier et al., 2019). Yet, we observed a delayed decrease in superoxide dismutase antioxidant activity when returning from the second foraging trip, and a slight increase in plasma non-enzymatic antioxidants. Both of these results combined with the lack of effect on oxidative damage may suggest a decrease in ROS production in response to CORT, as observed previously in lizards (Voituron et al., 2017). Indeed, lowered ROS production would decrease the need to use dietary acquired non-enzymatic antioxidants, thereby leading to an increase in their plasma levels, and could also decrease the need to produce endogenous antioxidants such as SOD. These results therefore suggest a potent resistance of king penguins to chronic glucocorticoid exposure in terms of oxidative stress. Such resistance mechanisms may be especially useful in long-lived animals exposed to a variety of long-term stressors, where reducing the physiological costs of stress will likely improve survival and lifetime reproductive success. However, it is important to note that our study focused solely on circulating markers, which may not have captured signs of oxidative damage in specific tissues such as the liver, muscle, or brain. In accordance, a meta-analysis related that among experimental studies involving GC implants, brain tissue was the most susceptible and heart tissue the least susceptible to GC-induced oxidative stress, while blood appeared to be moderately sensitive (Constantini et al., 2011).

We also evaluated the impact of chronic CORT increase on relative telomere length (rTL), a marker often used as a proxy of cellular ageing and fitness prospects (Wilbourn et al. 2018; Eastwood et al., 2019). Telomere shortening with age is known to be accelerated under both chronic stress and GC exposure (Angelier et al., 2017; Chatelain et al., 2019; Casagrande et al., 2023). While telomeres are not necessarily expected to shorten over a short experimental period (i.e. ∼45 days), recent work has suggested that telomeres may be regulatively shortened in response to energy imbalance and GCs through mTOR signalling (Casagrande & Hau, 2019). Relative telomere length did not differ significantly between treatment groups over the study period (p = 0.065), although the p-value approached the significance threshold. Given the absence of evidence for increased oxidative stress in CORT-implanted individuals (see above), this pattern is unlikely to reflect damage-induced telomere attrition (Reichert & Stier, 2017). Alternatively, it may reflect regulated telomere dynamics (Casagrande & Hau, 2019) or shifts in blood cell composition (i.e. changes in leukocyte proportions altering mean telomere length measured in whole blood without changes at the cellular level; Beaulieu et al., 2017). Investigating gene expression related to the mTOR pathway and telomere length in specific blood cell types would be needed to test these hypotheses.

Overall, our examination of physiological variables related to energy metabolism, oxidative stress and cellular ageing suggest that king penguins may be to some extent resilient and/or resistant to chronic GC exposure. Nonetheless, the sustained impairment of body condition observed in response to our experimental treatment suggests other costs to prolonged GC exposure. Since penguins and seabirds inherently have to contend with a dual environment, future studies are needed to understand if the decreased body condition we observed in CORT-treated individuals on-land may result from different performances when foraging at sea.

## Acknowledgments

We are grateful to the French Polar Institute (IPEV) and the Terres Australes et Antarctiques Françaises for providing financial and logistical support for this study through the polar program #119 (ECONERGY). This study is part of the long-term Studies in Ecology and Evolution (SEE-Life) program of the CNRS. We wish to thank the Zone Atelier Antarctique et Terres Australes (ZATA) from the CNRS for financial support, and the members of the Alfred Faure field station for their help and support in the field (especially Sylvain, Tom & Pierre). AS was supported by a ‘Turku Collegium for Science and Medicine’ Fellowship, a Marie Sklodowska-Curie Postdoctoral Fellowships (#894963) and the IdEx Université de Strasbourg (*HotPenguin*). SA and LA were funded by the French Polar Institute.

## Author contribution

Study design: AS, VAV, PBi. Funding acquisition: JPR, PBi, VAV, AS. Data collection in the field: AS, SA, TH, LA, and JPR. Data collection in the lab: ER and AS. Data analysis: AC, PBl, AS. Writing original draft: AC and AS. Writing review and editing: PBl, VAV, JPR, PBi, SA, TH, LA, ER.

## Ethical note

All the procedures were approved by the French Ethical Committee (APAFIS#16465-2018080111195526 v4) and the Terres Australes et Antarctiques Françaises (Arrêté TAAF A-2018-118).

## Declaration of no competing interests

The authors declare that they have no competing interests.

## Supplementary materials

**Fig. S1.**
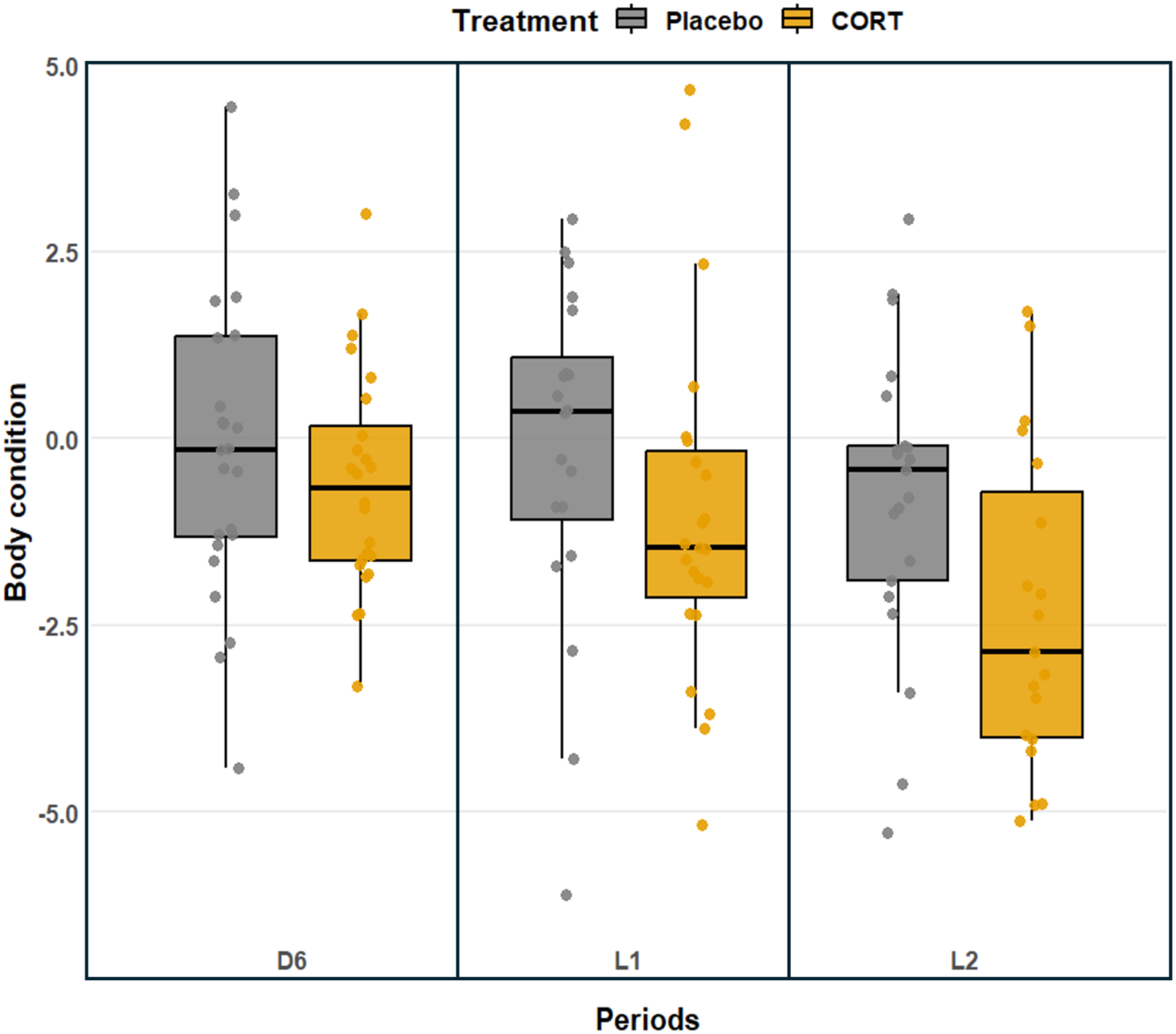
Boxplot of individual body condition between treatment across all periods. CORT-treated individuals show consistently lower body condition compared to Placebo (*F*_1,42.69_ = 6.15, p = 0.017). Treatment × Period did not show a significant interaction (p = 0.22), although a visual tendency of increasing difference between groups can be seen. Means are presented ± SE, * indicates p < 0.05, and N = 47, n = 129.

**Fig. S2.**
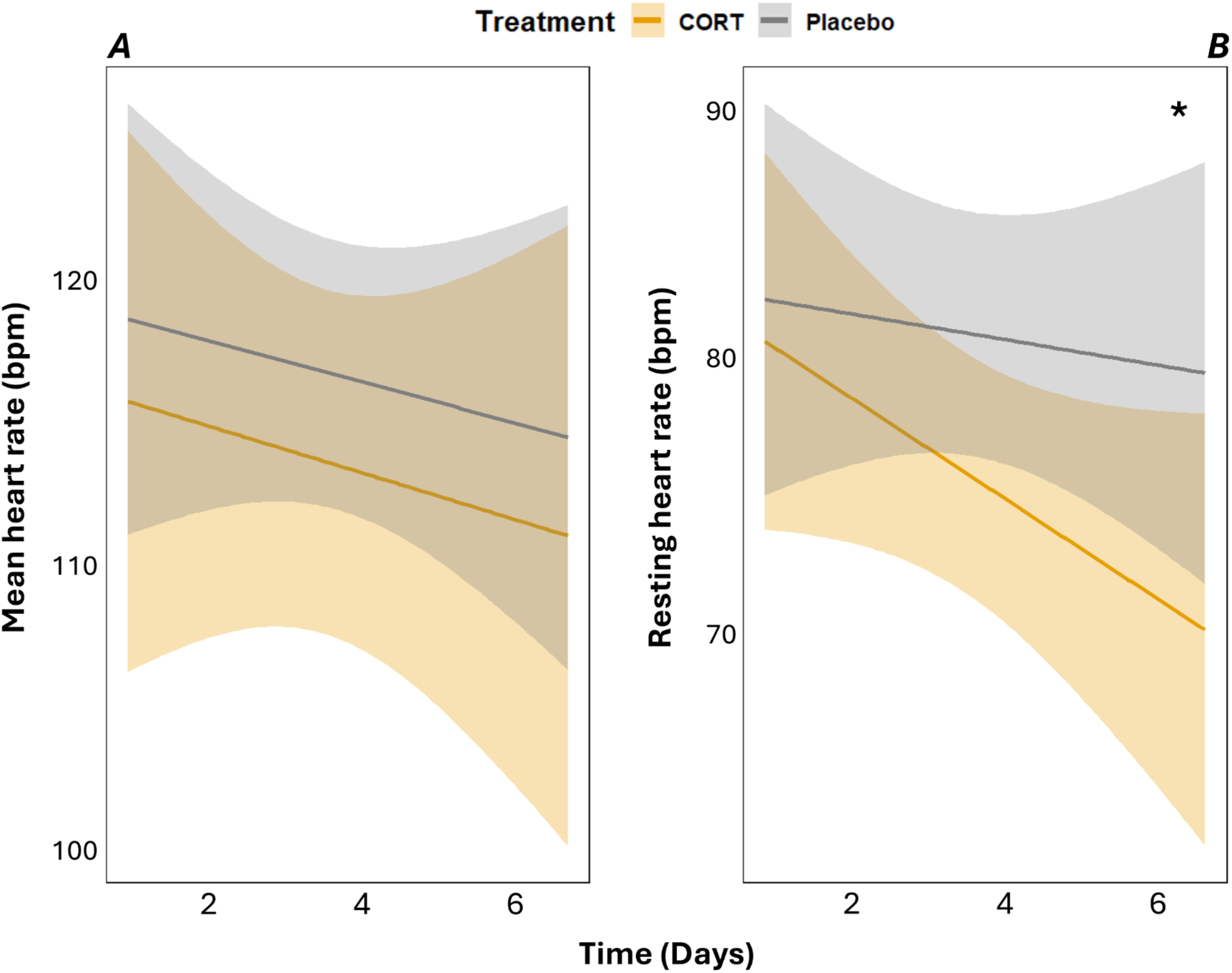
Effects of corticosterone treatment on mean heart rate (*A*) and resting heart rate (*B*) over time since implantation. Lines and points are coloured according to Treatment (yellow-orange for CORT; grey for Placebo). Shaded areas represent 95% confidence intervals. The interaction between time and Treatment is significant for resting heart rate, indicated by * on the graph (Time × Treatment: *F*_1,157.45_ = 5.13, p = 0.025) and N = 45, n = 203.

**Table S1A.**
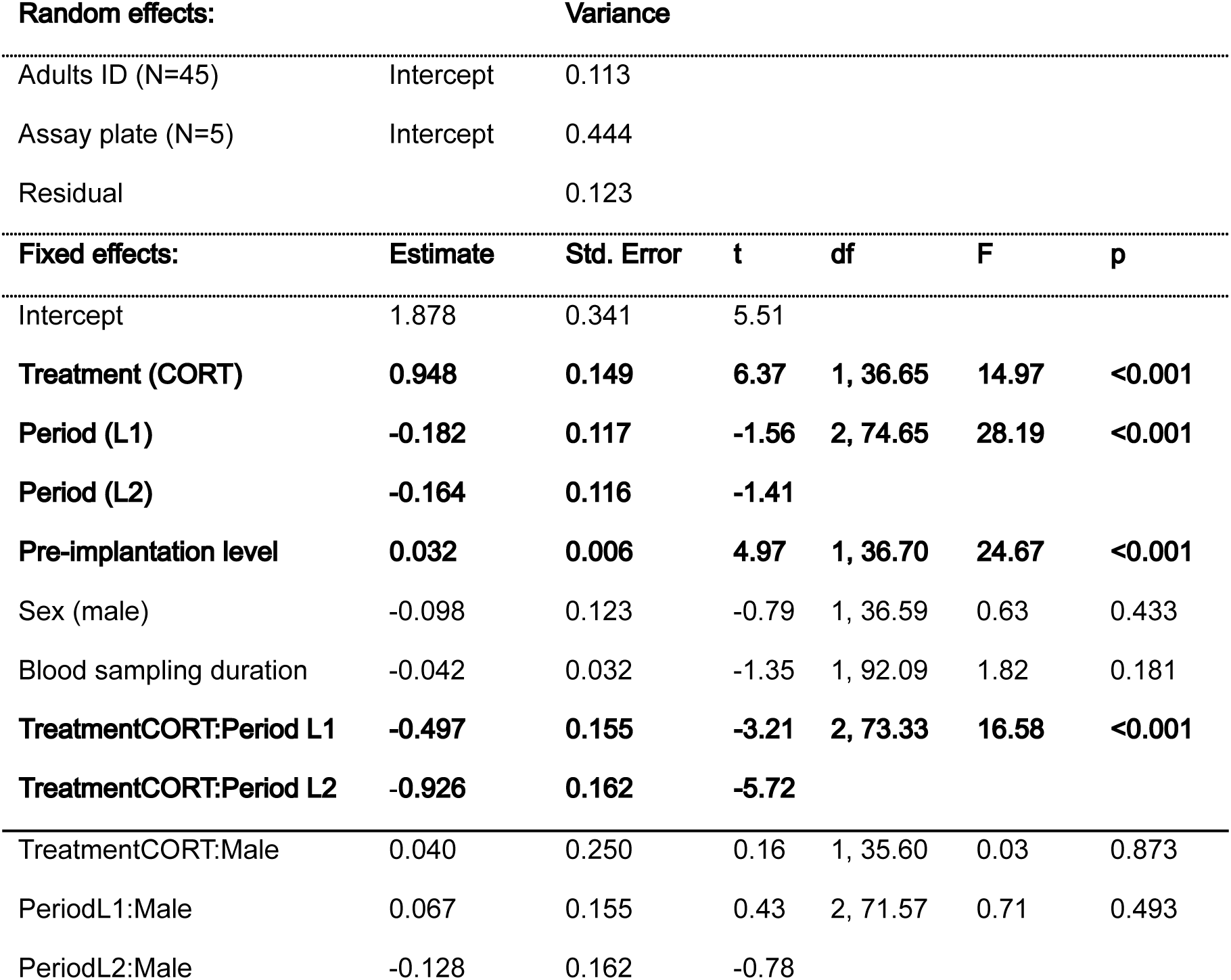
Log-transformed baseline corticosterone (n = 121). Random and fixed effects from the linear mixed model.

| Random effects: |  | Variance |  |  |  |  |
| --- | --- | --- | --- | --- | --- | --- |
| Adults ID (N=45) | Intercept | 0.113 |  |  |  |  |
| Assay plate (N=5) | Intercept | 0.444 |  |  |  |  |
| Residual |  | 0.123 |  |  |  |  |
| Fixed effects: | Estimate | Std. Error | t | df | F | p |
| Intercept | 1.878 | 0.341 | 5.51 |  |  |  |
| Treatment (CORT) | 0.948 | 0.149 | 6.37 | 1, 36.65 | 14.97 | <0.001 |
| Period (L1) | -0.182 | 0.117 | -1.56 | 2, 74.65 | 28.19 | <0.001 |
| Period (L2) | -0.164 | 0.116 | -1.41 |  |  |  |
| Pre-implantation level | 0.032 | 0.006 | 4.97 | 1, 36.70 | 24.67 | <0.001 |
| Sex (male) | -0.098 | 0.123 | -0.79 | 1, 36.59 | 0.63 | 0.433 |
| Blood sampling duration | -0.042 | 0.032 | -1.35 | 1, 92.09 | 1.82 | 0.181 |
| TreatmentCORT:Period L1 | -0.497 | 0.155 | -3.21 | 2, 73.33 | 16.58 | <0.001 |
| TreatmentCORT:Period L2 | -0.926 | 0.162 | -5.72 |  |  |  |
| TreatmentCORT:Male | 0.040 | 0.250 | 0.16 | 1, 35.60 | 0.03 | 0.873 |
| PeriodL1:Male | 0.067 | 0.155 | 0.43 | 2, 71.57 | 0.71 | 0.493 |
| PeriodL2:Male | -0.128 | 0.162 | -0.78 |  |  |  |

**Table S1B.**
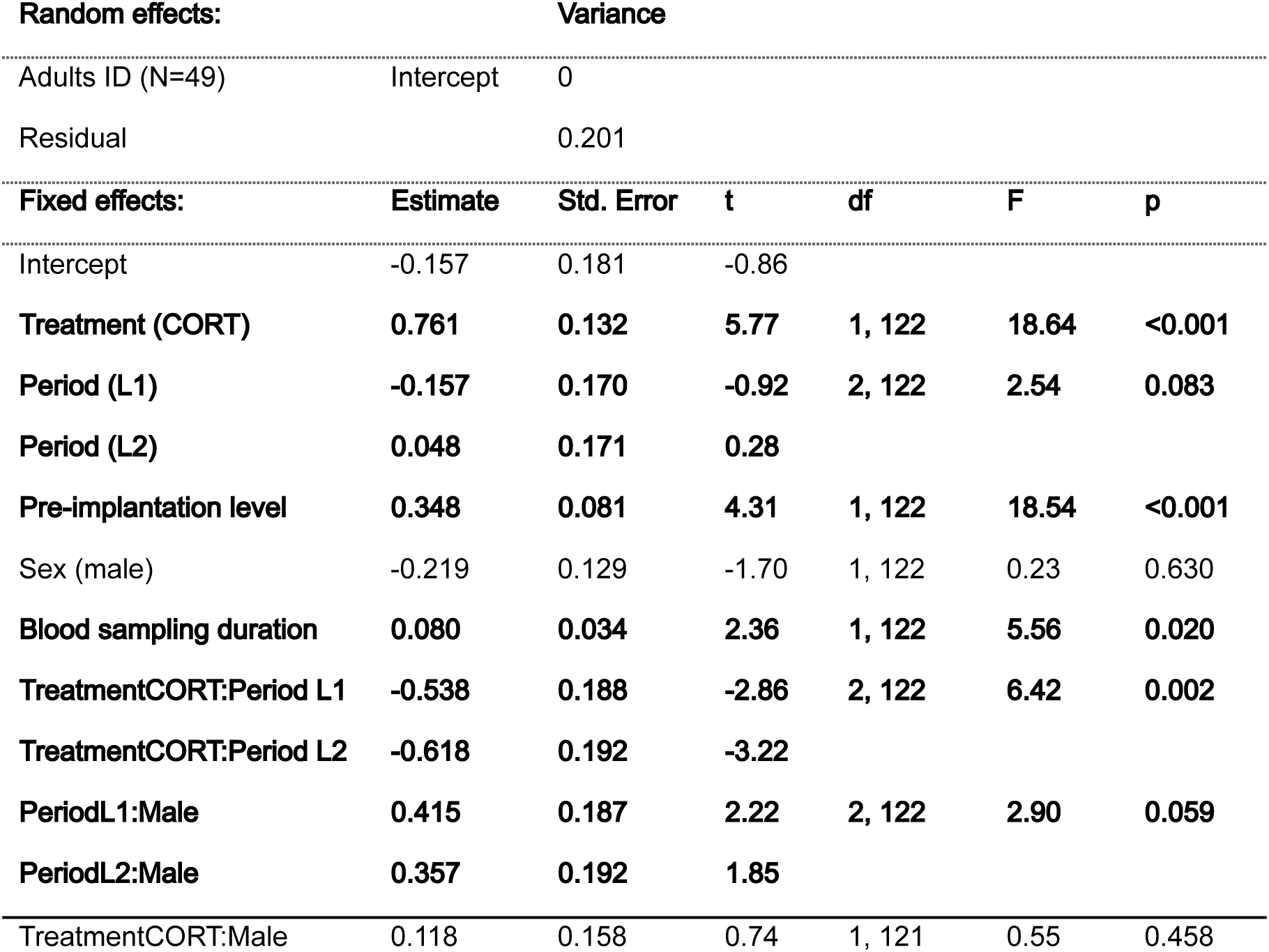
Log-transformed heterophyl-to-lymphocyte ratio (n = 133). Random and fixed effects from the linear mixed model.

**Table S2.** Body condition (n = 131). Random and fixed effects from the linear mixed model.

| **Random effects:** |  | **Variance** |  |  |  |  |
| --- | --- | --- | --- | --- | --- | --- |
| Adults ID (N=48) | Intercept | 0.86 |  |  |  |  |
| Experimenter (n=3) | Intercept | 0.02 |  |  |  |  |
| Residual |  | 2.56 |  |  |  |  |
| **Fixed effects:** | **Estimate** | **Std. Error** | **t** | **df** | **F** | **p** |
| Intercept | -0.941 | 0.470 | -2.00 |  |  |  |
| **Treatment (CORT)** | **-0.967** | **0.390** | **-2.48** | **1, 42.69** | **6.16** | **0.017** |
| **Period (L1)** | **-0.299** | **0.338** | **-0.89** | **2, 84.92** | **7.82** | **<0.001** |
| Period (L2) | -1.328 | 0.347 | -3.83 |  |  |  |
| Sex (male) | 0.068 | 0.394 | 0.17 | 1, 42.98 | 0.030 | 0.863 |
| **Pre-implantation level** | **0.482** | **0.098** | **4.93** | **1, 41.57** | **24.34** | **<0.001** |
| TreatmentCORT:Period L1 | -0.279 | 0.674 | -0.41 | 2, 83.11 | 1.52 | 0.224 |
| TreatmentCORT:Period L2 | -1.169 | 0.690 | -1.70 |  |  |  |
| TreatmentCORT:Male | -0.568 | 0.787 | -0.72 | 1, 41.76 | 0.52 | 0.47 |
| Period L1:Male | 0.165 | 0.683 | 0.24 | 2, 81.09 | 0.05 | 0.95 |
| Period L2:Male | 0.199 | 0.700 | 0.29 |  |  |  |

**Table S3A.**
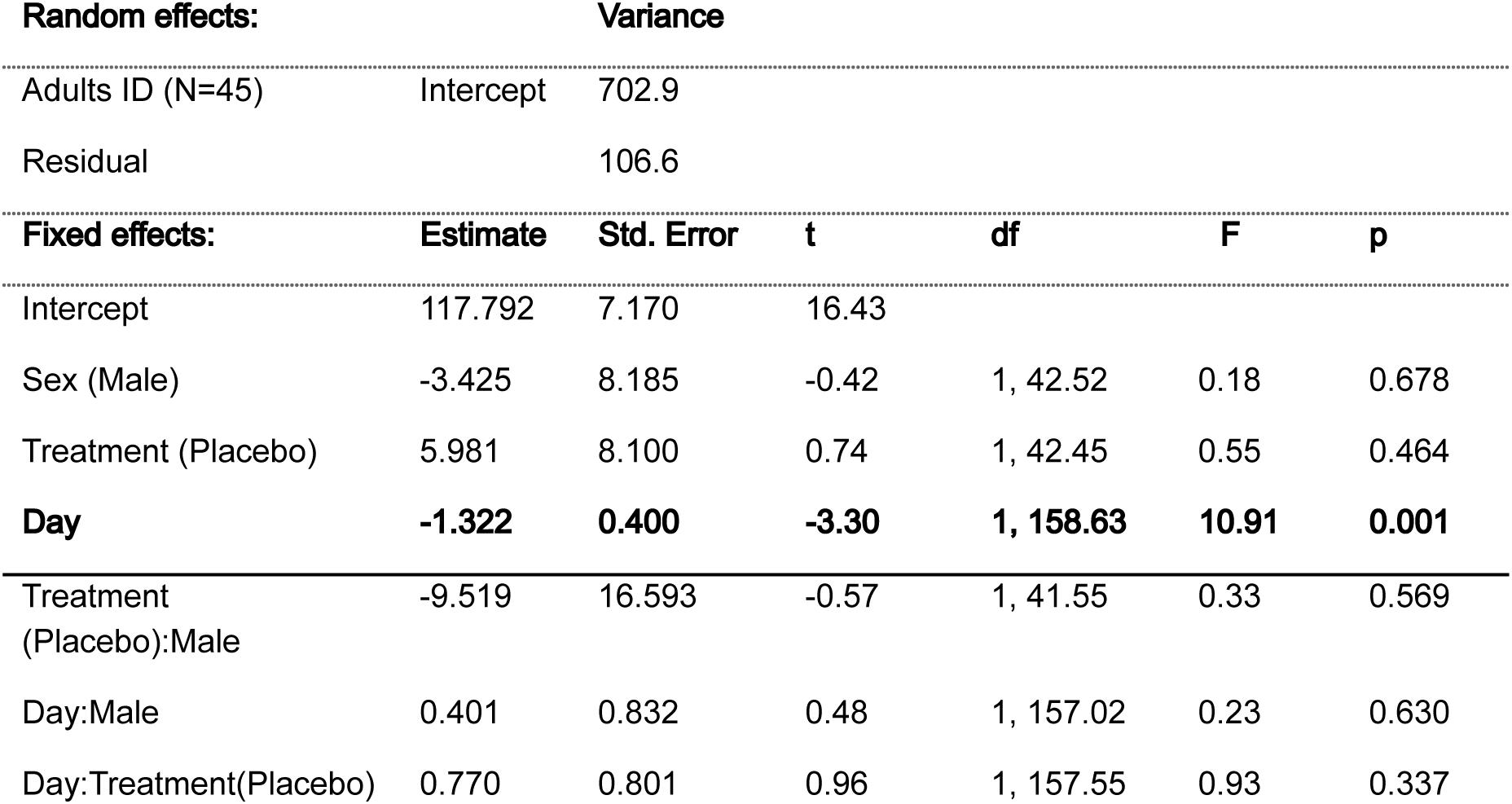
Mean heart rate (n = 203). Random and fixed effects from the linear mixed model.

**Table S3B.**
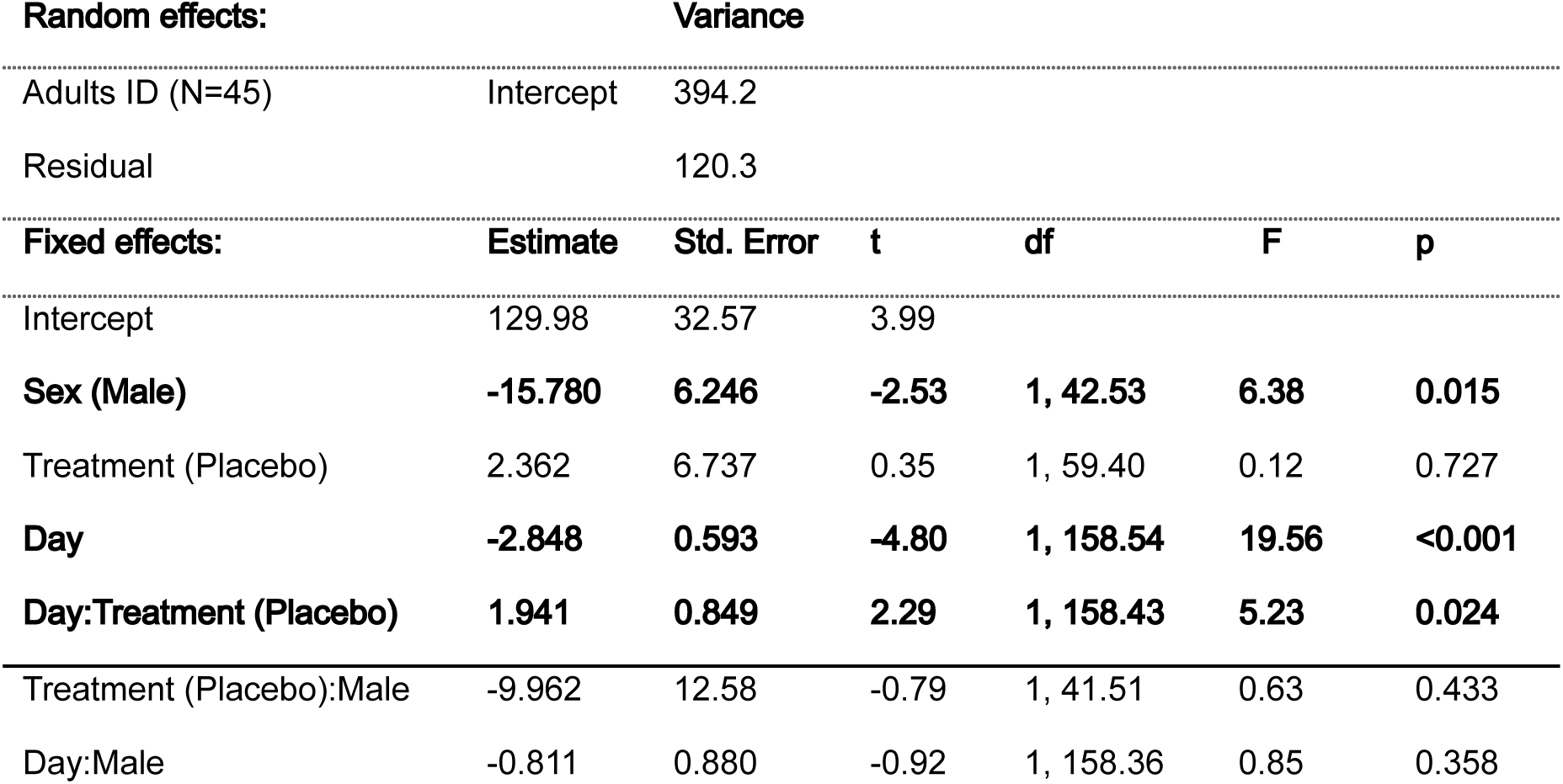
Resting heart rate (n = 203). Random and fixed effects from the linear mixed model.

**Table S4A.** Log-transformed glucose (n = 122). Random and fixed effects from the linear mixed model.

| **Random effects:** |  | **Variance** |  |  |  |  |
| --- | --- | --- | --- | --- | --- | --- |
| Adults ID (N=45) | Intercept | 0.005 |  |  |  |  |
| Assay plate (n=5) | Intercept | 0 |  |  |  |  |
| Residual |  | 0.020 |  |  |  |  |
| **Fixed effects:** | **Estimate** | **Std. Error** | **t** | **df** | **F** | **p** |
| Intercept | 0.130 | 0.116 | 1.12 |  |  |  |
| Treatment (CORT) | 0.098 | 0.048 | 2.07 | 1, 41.00 | 0.38 | 0.541 |
| Period (L1) | 0.132 | 0.046 | 2.84 | 2, 78.61 | 2.30 | 0.107 |
| Period (L2) | 0.104 | 0.046 | 2.24 |  |  |  |
| Sex (male) | 0.035 | 0.034 | 1.02 | 1, 38.93 | 1.03 | 0.315 |
| **Pre-implantation level** | **0.147** | **0.059** | **2.50** | **1, 44.70** | **6.26** | **0.016** |
| Blood sampling duration | 0.012 | 0.012 | 1.01 | 1, 109.00 | 1.00 | 0.319 |
| **TreatmentCORT: Period L1** | **-0.130** | **0.062** | **-2.10** | **2, 76.41** | **2.45** | **0.093** |
| **TreatmentCORT:Period L2** | **-0.101** | **0.064** | **-1.58** |  |  |  |
| TreatmentCORT:Male | 0.054 | 0.072 | 0.75 | 1, 38.61 | 0.560 | 0.459 |
| PeriodL1:Male | -0.033 | 0.061 | -0.54 | 2, 74.67 | 1.870 | 0.161 |
| PeriodL2:SexMale | -0.120 | 0.063 | -1.89 |  |  |  |

**Table S4B.**
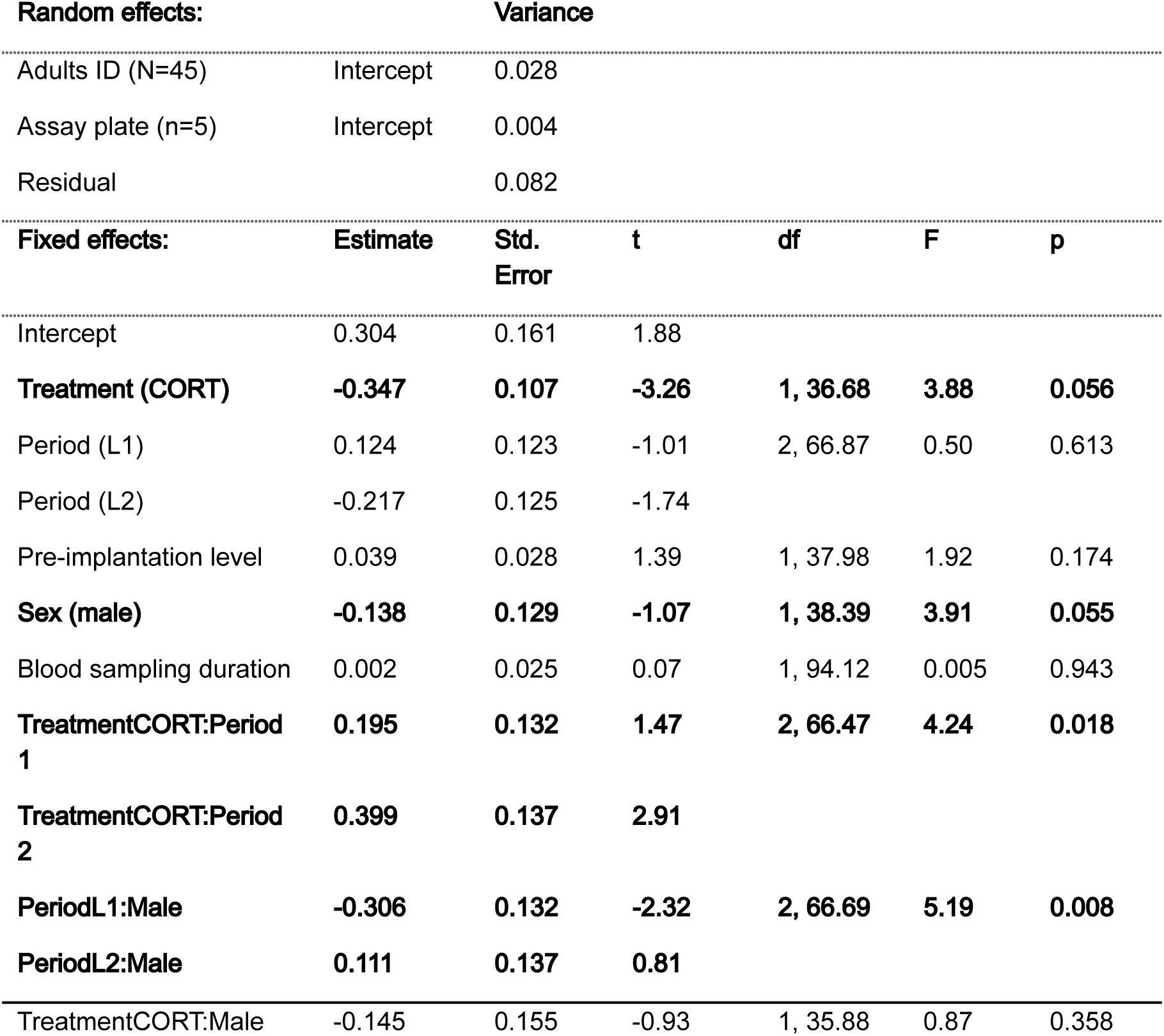
Log-transformed triglyceride (n = 114). Random and fixed effects from the linear mixed model.

**Table S4C.** Log-transformed uric acid (n = 114). Random and fixed effects from the linear mixed model.

| **Random effects:** |  | **Variance** |  |  |  |  |
| --- | --- | --- | --- | --- | --- | --- |
| Adults ID (N=45) | Intercept | 0.017 |  |  |  |  |
| Assay plate (n=5) | Intercept | 0.043 |  |  |  |  |
| Residual |  | 0.085 |  |  |  |  |
| **Fixed effects:** | **Estimate** | **Std. Error** | **t** | **df** | **F** | **p** |
| Intercept | -3.406 | 0.174 | -19.59 |  |  |  |
| **Treatment (CORT)** | **0.349** | **0.104** | **3.34** | **1, 36.01** | **3.33** | **0.076** |
| **Period (L1)** | **0.049** | **0.124** | **0.39** | **2, 72.03** | **5.93** | **0.004** |
| **Period (L2)** | **-0.189** | **0.127** | **-1.49** |  |  |  |
| Pre-implantation level | -1.101 | 1.344 | -0.82 | 1, 37.47 | 0.67 | 0.418 |
| Sex (male) | -0.193 | 0.114 | -1.69 | 1, 35.47 | 1.55 | 0.222 |
| Blood sampling duration | 0.023 | 0.025 | 0.90 | 1, 96.92 | 0.82 | 0.368 |
| **TreatmentCORT:Period L1** | **-0.270** | **0.134** | **-2.02** | **2, 71.38** | **4.19** | **0.019** |
| **TreatmentCORT:Period L2** | **-0.389** | **0.139** | **-2.79** |  |  |  |
| **PeriodL1:Male** | **-0.011** | **0.134** | **-0.08** | **2, 71.68** | **2.66** | **0.077** |
| **PeriodL2:Male** | **0.272** | **0.139** | **1.96** |  |  |  |
| TreatmentCORT:Male | -0.060 | 0.143 | -0.42 | 1, 35.32 | 0.18 | 0.675 |

**Table S4D.** Log-transformed lactate (n = 114). Random and fixed effects from the linear mixed model.

| **Random effects:** |  | **Variance** |  |  |  |  |
| --- | --- | --- | --- | --- | --- | --- |
| Adults ID (N=44) | Intercept | <0.0001 |  |  |  |  |
| Assay plate (n=5) | Intercept | 0.002 |  |  |  |  |
| Residual |  | 0.232 |  |  |  |  |
| **Fixed effects:** | **Estimate** | **Std. Error** | **t** | **df** | **F** | **p** |
| Intercept | -1.961 | 0.149 | -13.14 |  |  |  |
| Treatment (CORT) | -0.098 | 0.093 | -1.05 | 1, 104.95 | 1.11 | 0.295 |
| Period (L1) | 0.152 | 0.113 | 1.34 | 2, 102.27 | 2.21 | 0.115 |
| Period (L2) | 0.246 | 0.118 | 2.08 |  |  |  |
| Sex (male) | -0.128 | 0.098 | -1.31 | 1, 104.44 | 1.71 | 0.194 |
| Pre-implantation level | -0.487 | 0.465 | -1.05 | 1, 104.93 | 1.09 | 0.298 |
| **Blood sampling duration** | **0.159** | **0.0382** | **4.15** | **1, 104.86** | **17.24** | **<0.001** |
| TreatmentCORT:Male | -0.126 | 0.183 | -0.69 | 1, 37.35 | 0.47 | 0.496 |
| TreatmentCORT:PeriodL1 | 0.391 | 0.215 | 1.82 | 2, 98.77 | 1.78 | 0.174 |
| TreatmentCORT:PeriodL2 | 0.290 | 0.222 | 1.31 |  |  |  |
| PeriodL1:Male | -0.037 | 0.217 | -0.17 | 2, 101.33 | 2.29 | 0.106 |
| PeriodL2:Male | 0.402 | 0.223 | 1.80 |  |  |  |

**Table S5A.** Log-transformed plasma ROMs (n = 110). Random and fixed effects from the linear mixed model.

| Random effects: |  | Variance |  |  |  |  |
| --- | --- | --- | --- | --- | --- | --- |
| Adults ID (N=41) | Intercept | 0.037 |  |  |  |  |
| Assay plate (n=4) | Intercept | 0 |  |  |  |  |
| Residual |  | 0.534 |  |  |  |  |
| Fixed effects: | Estimate | Std. Error | t | df | F | p |
| Intercept | 0.672 | 0.263 | 2.56 |  |  |  |
| Treatment (CORT) | -0.277 | 0.225 | -1.23 | 1, 36.80 | 0.001 | 0.992 |
| Period (L1) | -0.227 | 0.174 | -1.31 | 2, 75.81 | 12.95 | <0.001 |
| Period (L2) | -0.901 | 0.185 | -4.89 |  |  |  |
| Pre-implantation level | 0.095 | 0.022 | 4.30 | 1, 37.56 | 18.47 | <0.001 |
| Sex (male) | -0.110 | 0.227 | -0.49 | 1, 35.74 | 1.07 | 0.307 |
| Blood sampling duration | 0.058 | 0.058 | 0.98 | 1, 101.94 | 0.97 | 0.327 |
| TreatmentCORT:Male | 0.551 | 0.306 | 1.80 | 1, 36.17 | 3.24 | 0.080 |
| TreatmentCORT:PeriodL1 | 0.312 | 0.335 | 0.93 | 2, 71.09 | 0.48 | 0.623 |
| TreatmentCORT:PeriodL2 | 0.242 | 0.352 | 0.69 |  |  |  |
| PeriodL1:Male | 0.080 | 0.339 | 0.24 | 2, 69.76 | 0.03 | 0.971 |
| PeriodL2:Male | 0.021 | 0.357 | 0.06 |  |  |  |

**Table S5B.** Log-transformed plasma 8-OHdG (n = 125). Random and fixed effects from the linear mixed model.

| **Random effects:** |  | **Variance** |  |  |  |  |
| --- | --- | --- | --- | --- | --- | --- |
| Adults ID (N=48) | Intercept | 0.070 |  |  |  |  |
| Assay plate (n=3) | Intercept | <0.001 |  |  |  |  |
| Residual |  | 0.096 |  |  |  |  |
| **Fixed effects:** | **Estimate** | **Std. Error** | **t** | **df** | **F** | **p** |
| Intercept | 3.602 | 0.124 | 29.00 |  |  |  |
| Treatment (CORT) | 0.229 | 0.125 | 1.84 | 1, 43.71 | 0.50 | 0.481 |
| Period (L1) | 0.106 | 0.099 | 1.07 | 2, 75.54 | 3.37 | 0.039 |
| Period (L2) | -0.092 | 0.097 | -0.94 |  |  |  |
| Sex (male) | -0.117 | 0.096 | -1.22 | 1, 42.37 | 1.50 | 0.227 |
| Blood sampling duration | -0.023 | 0.028 | -0.80 | 1, 101.37 | 0.65 | 0.424 |
| **TreatmentCORT:PeriodL1** | **-0.304** | **0.139** | **-2.20** | **2, 75.52** | **2.43** | **0.095** |
| **TreatmentCORT:PeriodL2** | **-0.178** | **0.141** | **-1.26** |  |  |  |
| TreatmentCORT:Male | -0.178 | 0.192 | -0.92 | 1, 41.53 | 0.86 | 0.360 |
| PeriodL1:Male | 0.009 | 0.140 | 0.06 | 2, 73.63 | 0.63 | 0.534 |
| PeriodL2:Male | 0.144 | 0.143 | 1.00 |  |  |  |

**Table S5C.** Log-transformed RBC 8-OHdG (n = 126). Random and fixed effects from the linear mixed model.

| **Random effects:** |  | **Variance** |  |  |  |  |
| --- | --- | --- | --- | --- | --- | --- |
| Adults ID (N=48) | Intercept | 0.0305 |  |  |  |  |
| Assay plate (n=3) | Intercept | 0.0189 |  |  |  |  |
| Residual |  | 0.0725 |  |  |  |  |
| **Fixed effects:** | **Estimate** | **Std. Error** | **t** | **df** | **F** | **p** |
| Intercept | 3.897 | 0.124 | 31.38 |  |  |  |
| Treatment (CORT) | 0.009 | 0.072 | 0.12 | 1, 46.58 | 0.01 | 0.902 |
| Period (L1) | 0.018 | 0.061 | 0.29 | 2, 83.30 | 0.51 | 0.602 |
| Period (L2) | -0.041 | 0.063 | -0.66 |  |  |  |
| Sex (male) | 0.058 | 0.070 | 0.83 | 1, 44.22 | 0.68 | 0.412 |
| Blood sampling duration | -0.019 | 0.023 | -0.82 | 1, 110.55 | 0.67 | 0.416 |
| TreatmentCORT:Male | 0.077 | 0.152 | 0.51 | 1, 47.73 | 0.26 | 0.613 |
| TreatmentCORT:PeriodL1 | -0.028 | 0.118 | -0.23 | 2, 80.80 | 1.33 | 0.271 |
| TreatmentCORT:PeriodL2 | -0.183 | 0.120 | -1.52 |  |  |  |
| PeriodL1:Male | -0.037 | 0.120 | -0.31 | 2, 79.34 | 0.05 | 0.954 |
| PeriodL2:Male | -0.021 | 0.123 | -0.17 |  |  |  |

**Table S5D.** RBC GPx (n = 115). Random and fixed effects from the linear mixed model.

| Random effects: |  | Variance |  |  |  |  |
| --- | --- | --- | --- | --- | --- | --- |
| Adults ID (N=45) | Intercept | 0.033 |  |  |  |  |
| Assay plate (n=5) | Intercept | 0.006 |  |  |  |  |
| Residual |  | 0.069 |  |  |  |  |
| Fixed effects: | Estimate | Std. Error | t | df | F | p |
| Intercept | 0.242 | 0.184 | 1.31 |  |  |  |
| Treatment (CORT) | -0.002 | 0.075 | -0.02 | 1, 37.63 | 0.01 | 0.982 |
| **BCA** | **0.046** | **0.007** | **6.77** | **1, 104.96** | **45.89** | **<0.001** |
| **Period (L1)** | **-0.185** | **0.064** | **-2.87** | **2, 77.15** | **4.51** | **0.014** |
| **Period (L2)** | **-0.159** | **0.070** | **-2.27** |  |  |  |
| **Pre-implantation level** | **0.418** | **0.082** | **5.08** | **1, 34.94** | **25.77** | **<0.001** |
| Sex (male) | -0.115 | 0.075 | -1.54 | 1, 37.97 | 2.36 | 0.133 |
| **Blood sampling duration** | **-0.070** | **0.023** | **-3.03** | **1, 101.25** | **9.18** | **0.003** |
| TreatmentCORT:Male | -0.171 | 0.152 | -1.12 | 1, 40.11 | 1.25 | 0.269 |
| TreatmentCORT:PeriodL1 | -0.138 | 0.122 | -1.13 | 2, 72.09 | 0.65 | 0.527 |
| TreatmentCORT:PeriodL2 | -0.073 | 0.125 | -0.59 |  |  |  |
| PeriodL1:Male | -0.084 | 0.121 | -0.69 | 2, 69.60 | 0.40 | 0.669 |
| PeriodL2:Male | -0.106 | 0.128 | -0.83 |  |  |  |

**Table S5E.**
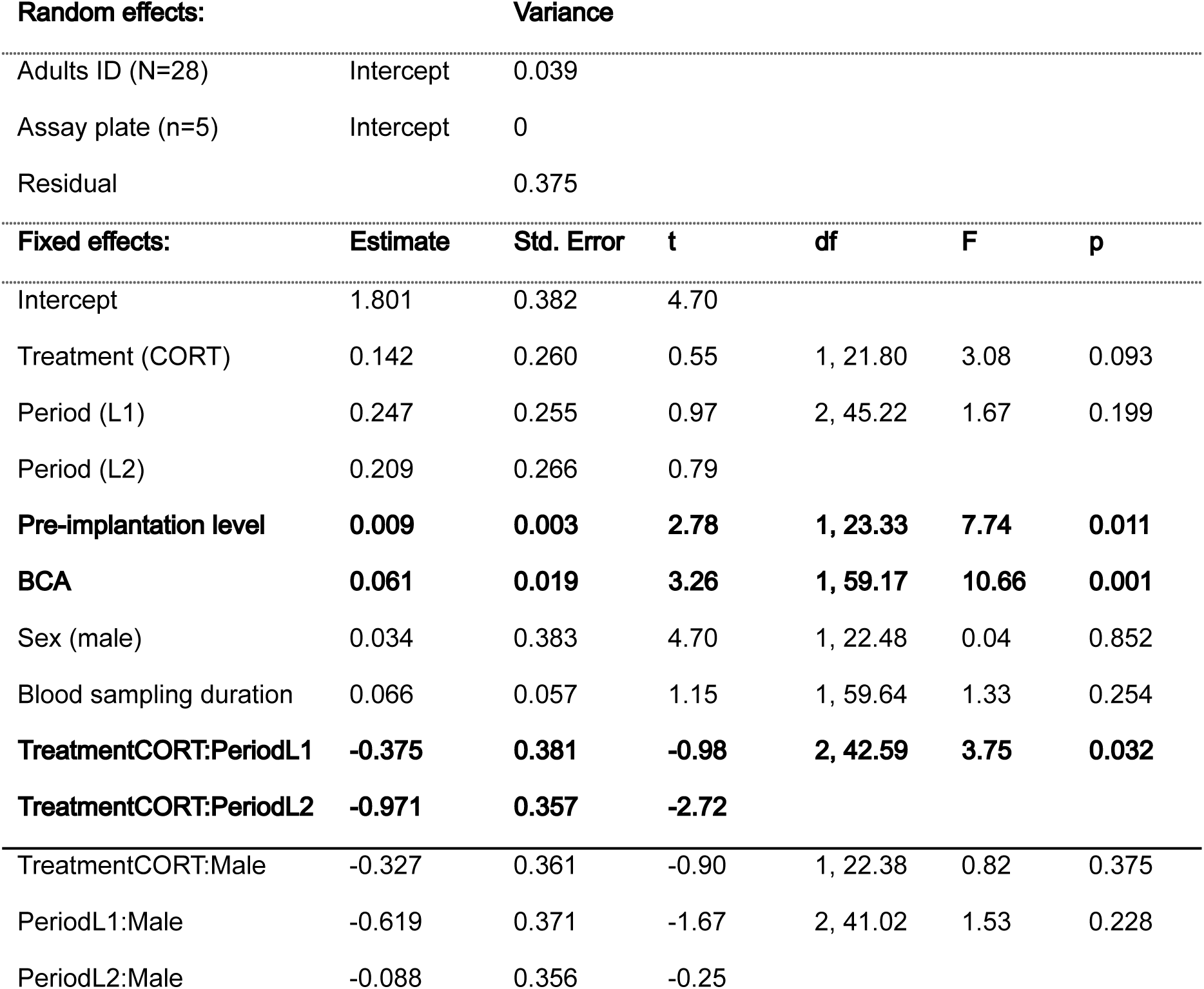
Log-transformed SOD (n = 70). Random and fixed effects from the linear mixed model.

| **Random effects:** |  | **Variance** |  |  |  |  |
| --- | --- | --- | --- | --- | --- | --- |
| Adults ID (N=28) | Intercept | 0.039 |  |  |  |  |
| Assay plate (n=5) | Intercept | 0 |  |  |  |  |
| Residual |  | 0.375 |  |  |  |  |
| **Fixed effects:** | **Estimate** | **Std. Error** | **t** | **df** | **F** | **p** |
| Intercept | 1.801 | 0.382 | 4.70 |  |  |  |
| Treatment (CORT) | 0.142 | 0.260 | 0.55 | 1, 21.80 | 3.08 | 0.093 |
| Period (L1) | 0.247 | 0.255 | 0.97 | 2, 45.22 | 1.67 | 0.199 |
| Period (L2) | 0.209 | 0.266 | 0.79 |  |  |  |
| **Pre-implantation level** | **0.009** | **0.003** | **2.78** | **1, 23.33** | **7.74** | **0.011** |
| **BCA** | **0.061** | **0.019** | **3.26** | **1, 59.17** | **10.66** | **0.001** |
| Sex (male) | 0.034 | 0.383 | 4.70 | 1, 22.48 | 0.04 | 0.852 |
| Blood sampling duration | 0.066 | 0.057 | 1.15 | 1, 59.64 | 1.33 | 0.254 |
| **TreatmentCORT:PeriodL1** | **-0.375** | **0.381** | **-0.98** | **2, 42.59** | **3.75** | **0.032** |
| **TreatmentCORT:PeriodL2** | **-0.971** | **0.357** | **-2.72** |  |  |  |
| TreatmentCORT:Male | -0.327 | 0.361 | -0.90 | 1, 22.38 | 0.82 | 0.375 |
| PeriodL1:Male | -0.619 | 0.371 | -1.67 | 2, 41.02 | 1.53 | 0.228 |
| PeriodL2:Male | -0.088 | 0.356 | -0.25 |  |  |  |

**Table S5F.**
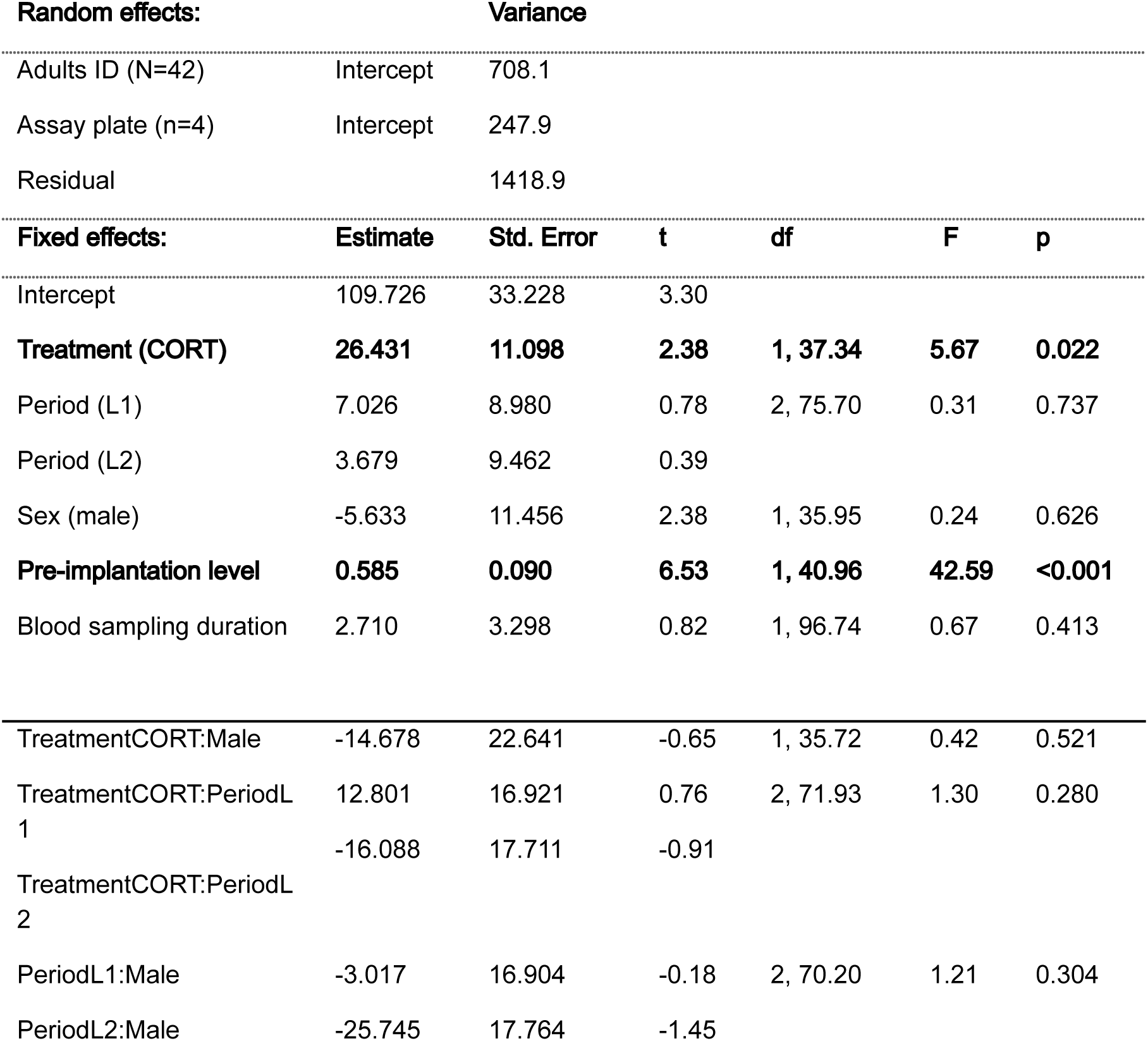
Plasma OXY (n = 114). Random and fixed effects from the linear mixed model.

| **Random effects:** |  | **Variance** |  |  |  |  |
| --- | --- | --- | --- | --- | --- | --- |
| Adults ID (N=42) | Intercept | 708.1 |  |  |  |  |
| Assay plate (n=4) | Intercept | 247.9 |  |  |  |  |
| Residual |  | 1418.9 |  |  |  |  |
| **Fixed effects:** | **Estimate** | **Std. Error** | **t** | **df** | **F** | **p** |
| Intercept | 109.726 | 33.228 | 3.30 |  |  |  |
| **Treatment (CORT)** | **26.431** | **11.098** | **2.38** | **1, 37.34** | **5.67** | **0.022** |
| Period (L1) | 7.026 | 8.980 | 0.78 | 2, 75.70 | 0.31 | 0.737 |
| Period (L2) | 3.679 | 9.462 | 0.39 |  |  |  |
| Sex (male) | -5.633 | 11.456 | 2.38 | 1, 35.95 | 0.24 | 0.626 |
| **Pre-implantation level** | **0.585** | **0.090** | **6.53** | **1, 40.96** | **42.59** | **<0.001** |
| Blood sampling duration | 2.710 | 3.298 | 0.82 | 1, 96.74 | 0.67 | 0.413 |
| TreatmentCORT:Male | -14.678 | 22.641 | -0.65 | 1, 35.72 | 0.42 | 0.521 |
| TreatmentCORT:PeriodL1 | 12.801 | 16.921 | 0.76 | 2, 71.93 | 1.30 | 0.280 |
| TreatmentCORT:PeriodL2 | -16.088 | 17.711 | -0.91 |  |  |  |
| PeriodL1:Male | -3.017 | 16.904 | -0.18 | 2, 70.20 | 1.21 | 0.304 |
| PeriodL2:Male | -25.745 | 17.764 | -1.45 |  |  |  |

**Table S5G.**
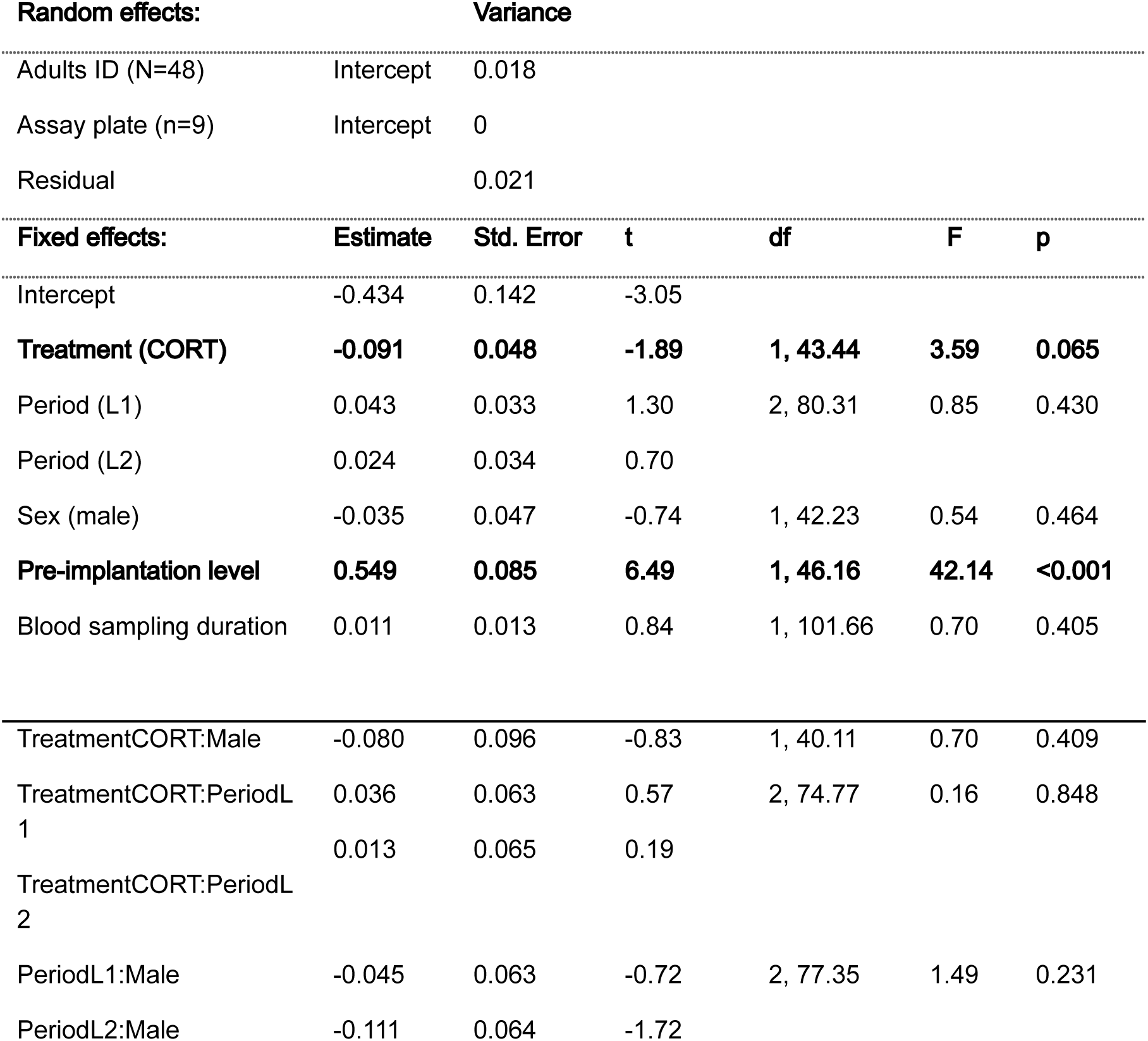
Log-transformed rTL (n = 128). Random and fixed effects from the linear mixed model.

## Notes

### Competing Interest Statement

The authors have declared no competing interest.

## References

Angelier, F., Shaffer, S. A., Weimerskirch, H., Trouvé, C., & Chastel, O. (2007). Corticosterone and Foraging Behavior in a Pelagic Seabird. Physiological and Biochemical Zoology, 80(3), 283–292. 10.1086/512585

Angelier, F., Bost, C.-A., Giraudeau, M., Bouteloup, G., Dano, S., & Chastel, O. (2008). Corticosterone and foraging behavior in a diving seabird: The Adélie penguin, Pygoscelis adeliae. General and Comparative Endocrinology, 156(1), 134–144. 10.1016/j.ygcen.2007.12.001

Angelier, F., Clément-Chastel, C., Welcker, J., Gabrielsen, G. W., & Chastel, O. (2009). How does corticosterone affect parental behaviour and reproductive success? A study of prolactin in black-legged kittiwakes. Functional Ecology, 23(4), 784–793. 10.1111/j.1365-2435.2009.01545.x

Angelier, F., Wingfield, J. C., Weimerskirch, H., & Chastel, O. (2010). Hormonal correlates of individual quality in a long-lived bird: A test of the ‘corticosterone–fitness hypothesis’. Biology Letters, 6(6), 846–849. 10.1098/rsbl.2010.0376

Angelier, F., & Wingfield, J. C. (2013). Importance of the glucocorticoid stress response in a changing world: Theory, hypotheses and perspectives. General and Comparative Endocrinology, 190, 118–128. 10.1016/j.ygcen.2013.05.022

Angelier, F., Costantini, D., Blévin, P. and Chastel, O. (2017). Do glucocorticoids mediate the link between environmental conditions and telomere dynamics in wild vertebrates? A review. General and comparative endocrinology 1–13. 10.1016/j.ygcen.2017.07.007

Barbraud, C., Delord, K., Bost, C. A., Chaigne, A., Marteau, C., & Weimerskirch, H. (2020). Population trends of penguins in the French Southern Territories. Polar Biology, 43(7), 835–850. 10.1007/s00300-020-02691-6

Beaulieu, M., Benoit, L., Abaga, S., Kappeler, P. M., & Charpentier, M. J. E. (2017). Mind the cell: Seasonal variation in telomere length mirrors changes in leucocyte profile. Molecular Ecology, 26(20), 5603–5613. 10.1111/mec.14329

Belthoff, J. R., & Dufty, J., Alfred M. (1998). Corticosterone, body condition and locomotor activity: A model for dispersal in screech-owls. Animal Behaviour, 55(2), 405–415. 10.1006/anbe.1997.0625

Bernard, S. F., Orvoine, J., & Groscolas, R. (2003). Glucose regulates lipid metabolism in fasting king penguins. American Journal of Physiology-Regulatory, Integrative and Comparative Physiology, 285(2), R313–R320. 10.1152/ajpregu.00094.2003

Bize, P., Stocker, A., Jenni-Eiermann, S., Gasparini, J., & Roulin, A. (2010). Sudden weather deterioration but not brood size affects baseline corticosterone levels in nestling Alpine swifts. Hormones and Behavior, 58(4), 591–598. 10.1016/j.yhbeh.2010.06.020

Blackburn, E. H. (2000). Telomeres and telomerase. The Keio journal of medicine, 49(2), 59–65.

Blas, J., Baos, R., Bortolotti, G. R., Marchant, T. A., & Hiraldo, F. (2006). Age-related variation in the adrenocortical response to stress in nestling white storks (Ciconia ciconia) supports the developmental hypothesis. General and Comparative Endocrinology, 148(2), 172–180. 10.1016/j.ygcen.2006.02.011

Bonier, F., Martin, P. R., Moore, I. T., & Wingfield, J. C. (2009). Do baseline glucocorticoids predict fitness? Trends in Ecology & Evolution, 24(11), 634–642. 10.1016/j.tree.2009.04.013

Braun, E. J., & Sweazea, K. L. (2008). Glucose regulation in birds. Comparative Biochemistry and Physiology Part B: Biochemistry and Molecular Biology, 151(1), 1–9. 10.1016/j.cbpb.2008.05.007

Breuner, C. W., Patterson, S. H., & Hahn, T. P. (2008). In search of relationships between the acute adrenocortical response and fitness. General and Comparative Endocrinology, 157(3), 288–295. 10.1016/j.ygcen.2008.05.017

Breuner, C. W., Delehanty, B., & Boonstra, R. (2013). Evaluating stress in natural populations of vertebrates: Total CORT is not good enough. Functional Ecology, 27(1), 24–36. 10.1111/1365-2435.12016

Brisson-Curadeau, É., Bost, C.-A., Cherel, Y., & Elliott, K. (2024). King Penguins adjust foraging effort rather than diet when faced with poor foraging conditions. Ibis, 166(2), 723–731. 10.1111/ibi.13287

Butler, M. W., Armour, E. M., Minnick, J. A., Rossi, M. L., Schock, S. F., Berger, S. E., & Hines, J. K. (2020). Effects of stress-induced increases of corticosterone on circulating triglyceride levels, biliverdin concentration, and heme oxygenase expression. Comparative Biochemistry and Physiology Part A: Molecular & Integrative Physiology, 240, 110608. 10.1016/j.cbpa.2019.110608

Cabezas, S., Blas, J., Marchant, T. A., & Moreno, S. (2007). Physiological stress levels predict survival probabilities in wild rabbits. Hormones and Behavior, 51(3), 313–320. 10.1016/j.yhbeh.2006.11.004

Cain, D. W., & Cidlowski, J. A. (2017). Immune regulation by glucocorticoids. Nature Reviews Immunology, 17(4), 233–247. 10.1038/nri.2017.1

Campbell, J. E., Peckett, A. J., D’souza, A. M., Hawke, T. J., & Riddell, M. C. (2011). Adipogenic and lipolytic effects of chronic glucocorticoid exposure. American Journal of Physiology-Cell Physiology, 300(1), C198–C209. 10.1152/ajpcell.00045.2010

Cannon, W. B. (1915). Bodily changes in pain, hunger, fear and rage: An account of recent researches into the function of emotional excitement. D. Appleton and Company. 10.1126/science.42.1089.696.b

Casagrande, S., & Hau, M. (2019). Telomere attrition: Metabolic regulation and signalling function? Biology Letters, 15(3), 20180885. 10.1098/rsbl.2018.0885

Casagrande, S., Loveland, J. L., Oefele, M., Boner, W., Lupi, S., Stier, A., & Hau, M. (2023). Dietary nucleotides can prevent glucocorticoid-induced telomere attrition in a fast-growing wild vertebrate. Molecular Ecology, 32(19), 5429–5447. 10.1111/mec.17114

Chatelain, M., Drobniak, S. M. and Szulkin, M. (2019). The association between stressors and telomeres in non-human vertebrates: a meta-analysis. Ecology Letters 99, 21–18. 10.1111/ele.13426

Cherel, Y., Robin, J. P., Walch, O., Karmann, H., Netchitailo, P., & Le Maho, Y. (1988). Fasting in king penguin. I. Hormonal and metabolic changes during breeding. The American Journal of Physiology, 254(2 Pt 2), R170–177. 10.1152/ajpregu.1988.254.2.R170

Clinchy, M., Sheriff, M. J., & Zanette, L. Y. (2013). Predator-induced stress and the ecology of fear. Functional Ecology, 27(1), 56–65. 10.1111/1365-2435.12007

Costantini, D., Fanfani, A., & Dell’omo, G. (2008). Effects of corticosteroids on oxidative damage and circulating carotenoids in captive adult kestrels (Falco tinnunculus). Journal of Comparative Physiology. B, Biochemical, Systemic, and Environmental Physiology, 178(7), 829–835. 10.1007/s00360-008-0270-z

Costantini, D., Marasco, V., & Møller, A. P. (2011). A meta-analysis of glucocorticoids as modulators of oxidative stress in vertebrates. Journal of Comparative Physiology B, 181(4), 447–456. 10.1007/s00360-011-0566-2

Costantini, D. (2016). Oxidative stress ecology and the d-ROMs test: Facts, misfacts and an appraisal of a decade’s work. Behavioral Ecology and Sociobiology, 70(5), 809–820. 10.1007/s00265-016-2091-5

Côté, S. D. (2000). Aggressiveness in king penguins in relation to reproductive status and territory location. Animal Behaviour, 59(4), 813–821. 10.1006/anbe.1999.1384

Cyr, N. E., & Romero, L. M. (2009). Identifying hormonal habituation in field studies of stress. General and Comparative Endocrinology, 161(3), 295–303. 10.1016/j.ygcen.2009.02.001

Dallman, M. F., Akana, S. F., Scribner, K. A., Bradbury, M. J., Walker, C. D., Strack, A. M., & Cascio, C. S. (1992). Stress, feedback and facilitation in the hypothalamo-pituitary-adrenal axis. Journal of neuroendocrinology, 4(5), 517–526. 10.1111/j.1365-2826.1992.tb00200.x

Davis, A. K., Maney, D. L., & Maerz, J. C. (2008). The use of leukocyte profiles to measure stress in vertebrates: A review for ecologists. Functional Ecology, 22(5), 760–772. 10.1111/j.1365-2435.2008.01467.x

Dhabhar, F. S. (2002). Stress-induced augmentation of immune function—The role of stress hormones, leukocyte trafficking, and cytokines. Brain, Behavior, and Immunity, 16(6), 785–798. 10.1016/S0889-1591(02)00036-3

Eastwood, J. R., Hall, M. L., Teunissen, N., Kingma, S. A., Aranzamendi, N. H., Fan, M., Roast, M., Verhulst, S. and Peters, A. (2019). Early-life telomere length predicts lifespan and lifetime reproductive success in a wild bird. Molecular Ecology 28, 1127–1137. 10.1111/mec.15002

Finkel, T., & Holbrook, N. J. (2000). Oxidants, oxidative stress and the biology of ageing. Nature, 408(6809), 239–247. 10.1038/35041687

Fox, J., & Weisberg, S. (2019). Using car functions in other functions. CRAN R.

Goldstein, D. S. (2010). Adrenal Responses to Stress. Cellular and Molecular Neurobiology, 30(8), 1433–1440. 10.1007/s10571-010-9606-9

Gouin, J.-P. (2011). Chronic Stress, Immune Dysregulation, and Health. American Journal of Lifestyle Medicine, 5(6), 476–485. 10.1177/1559827610395467

Groscolas, R., & Robin, J.-P. (2001). Long-term fasting and re-feeding in penguins. Comparative Biochemistry and Physiology Part A: Molecular & Integrative Physiology, The Physiological Consequences of Feeding in Animals, 128(3), 643–653. 10.1016/S1095-6433(00)00341-X

Groscolas, R., Lacroix, A., & Robin, J.-P. (2008). Spontaneous egg or chick abandonment in energy-depleted king penguins: A role for corticosterone and prolactin? Hormones and Behavior, 53(1), 51–60.

Groscolas, R., Viera, V., Guerin, N., Handrich, Y., & Côté, S. D. (2010). Heart rate as a predictor of energy expenditure in undisturbed fasting and incubating penguins. The Journal of Experimental Biology, 213(1), 153–160. 10.1242/jeb.033720

Gross, W. B., & Siegel, H. S. (1983). Evaluation of the heterophil/lymphocyte ratio as a measure of stress in chickens. Avian Diseases, 972–979. 10.2307/1590198

Handrich, Y., Bevan, R. M., Charrassin, J.-B., Butler, P. J., Ptz, K., Woakes, A. J., Lage, J., & Maho, Y. L. (1997). Hypothermia in foraging king penguins. Nature, 388(6637), 64–67. 10.1038/40392

Hartig, F. (2022, March). DHARMa: residual diagnostics for hierarchical (multi-level/mixed) regression models. R package version 0.4. 6.

Haussmann, M. F., & Marchetto, N. M. (2010). Telomeres: Linking stress and survival, ecology and evolution. Current Zoology, 56(6), 714–727. 10.1093/czoolo/56.6.714

Henderson, L. J., Evans, N. P., Heidinger, B. J., Herborn, K. A., & Arnold, K. E. (2017). Do glucocorticoids predict fitness?: Linking environmental conditions, corticosterone and reproductive success in the blue tit, Cyanistes caeruleus. Royal Society Open Science, 4(10), 170875. 10.1098/rsos.170875

Jaatinen, K., Seltmann, M. W., & Öst, M. (2014). Context-dependent stress responses and their connections to fitness in a landscape of fear. Journal of Zoology, 294(3), 147–153. 10.1111/jzo.12169

Jimeno, B., Hau, M., & Verhulst, S. (2018). Corticosterone levels reflect variation in metabolic rate, independent of ‘stress’. Scientific Reports, 8(1), 13020. 10.1038/s41598-018-31258-z

Kärkkäinen, T., Briga, M., Laaksonen, T., & Stier, A. (2022). Within-individual repeatability in telomere length: A meta-analysis in nonmammalian vertebrates. Molecular Ecology, 31(23), 6339–6359. 10.1111/mec.16155

Kitaysky, A. S., Kitaiskaia, E. V., Piatt, J. F., & Wingfield, J. C. (2003). Benefits and costs of increased levels of corticosterone in seabird chicks. Hormones and Behavior, 43(1), 140–149. 10.1016/S0018-506X(02)00030-2

Kitaysky, A. S., Piatt, J. F., & Wingfield, J. C. (2007). Stress hormones link food availability and population processes in seabirds. Marine Ecology Progress Series, 352, 245–258. 10.3354/meps07074

Kuznetsova, A., Brockhoff, P. B., & Christensen, R. H. B. (2017). lmerTest Package: Tests in Linear Mixed Effects Models. Journal of Statistical Software, 82, 1–26. 10.18637/jss.v082.i13

Landys, M. M., Ramenofsky, M., & Wingfield, J. C. (2006). Actions of glucocorticoids at a seasonal baseline as compared to stress-related levels in the regulation of periodic life processes. General and Comparative Endocrinology, 148(2), 132–149. 10.1016/j.ygcen.2006.02.013

Lane, S. J., Fossett, T. E. E., VanDiest, I. J., & Sewall, K. B. B. (2025). Recovery through resistance? Nesting urban female song sparrows (*Melospiza melodia*) have a lower glucocorticoid response to disturbance and return to parental care as quickly as rural females. Frontiers in Physiology, 16. 10.3389/fphys.2025.1520208

Le, P. P., Friedman, J. R., Schug, J., Brestelli, J. E., Parker, J. B., Bochkis, I. M., & Kaestner, K. H. (2005). Glucocorticoid Receptor-Dependent Gene Regulatory Networks. PLOS Genetics, 1(2), e16. 10.1371/journal.pgen.0010016

Le Maho, Y., Karmann, H., Briot, D., Handrich, Y., Robin, J. P., Mioskowski, E., Cherel, Y., & Farni, J. (1992). Stress in birds due to routine handling and a technique to avoid it. American Journal of Physiology-Regulatory, Integrative and Comparative Physiology, 263(4), R775–R781. 10.1152/ajpregu.1992.263.4.R775

Lemonnier, C., Schull, Q., Stier, A., Boonstra, R., Delehanty, B., Lefol, E., Durand, L., Pardonnet, S., Robin, J.-P., Criscuolo, F., Bize, P., & Viblanc, V. A. (2024). Social, not genetic, programming of development and stress physiology of a colonial seabird. Proceedings of the Royal Society B: Biological Sciences, 291(2028), 20240853. 10.1098/rspb.2024.0853

Lemonnier, C., Bost, C.-A., Joly, N., Stier, A., Robin, J.-P., Handrich, Y., Cillard, A., Montblanc, M., Bize, P., & Viblanc, V. A. (2025). Foraging flexibility in response to at-sea constraints in a deep diver, the king penguin: An experimental study. Oecologia, 207(7), 123. 10.1007/s00442-025-05754-9

Lemonnier, C., Colominas-Ciuró, R., Stier, A., Avril, S., Cillard, A., Dumas-Roussel, A., Meymy, C., Montblanc, M., Bost, C.-A., & Viblanc, V. A. (2026). Evaluating the links between heterophil-to-lymphocyte ratio, foraging effort and breeding success in two penguin species. Marine Biology, 173(2), 23. 10.1007/s00227-025-04763-9

Lenth, R., Singmann, H., Love, J., Buerkner, P., & Herve, M. (2019). Package ‘emmeans’. R package version, 1(3.2).

Lewden, A., Ward, C., Noiret, A., Avril, S., Abolivier, L., Gérard, C., Hammer, T. L., Raymond, É., Robin, J.-P., Viblanc, V. A., Bize, P., & Stier, A. (2024). Surface temperatures are influenced by handling stress independently of corticosterone levels in wild king penguins (Aptenodytes patagonicus). Journal of Thermal Biology, 121, 103850. 10.1016/j.jtherbio.2024.103850

Lin, H., Decuypere, E., & Buyse, J. (2004). Oxidative stress induced by corticosterone administration in broiler chickens (Gallus gallus domesticus) 1. Chronic exposure. Comparative Biochemistry and Physiology. Part B, Biochemistry & Molecular Biology, 139(4), 737–744. 10.1016/j.cbpc.2004.09.013

Lynn, S. E., Breuner, C. W., & Wingfield, J. C. (2003). Short-term fasting affects locomotor activity, corticosterone, and corticosterone binding globulin in a migratory songbird. Hormones and Behavior, 43(1), 150–157. 10.1016/S0018-506X(02)00023-5

MacDougall-Shackleton, S. A., Bonier, F., Romero, L. M., & Moore, I. T. (2019). Glucocorticoids and “Stress” Are Not Synonymous. Integrative Organismal Biology, 1(1), obz017. 10.1093/iob/obz017

Magomedova, L., & Cummins, C. L. (2016). Glucocorticoids and Metabolic Control. Handbook of Experimental Pharmacology, 233, 73–93. 10.1007/164_2015_1

Majer, A. D., Fasanello, V. J., Tindle, K., Frenz, B. J., Ziur, A. D., Fischer, C. P., Fletcher, K. L., Seecof, O. M., Gronsky, S., Vassallo, B. G., Reed, W. L., Paitz, R. T., Stier, A., & Haussmann, M. F. (2019). Is there an oxidative cost of acute stress? Characterization, implication of glucocorticoids and modulation by prior stress experience. Proceedings of the Royal Society B: Biological Sciences, 286(1915), 20191698. 10.1098/rspb.2019.1698

Manoli, I., Alesci, S., Blackman, M. R., Su, Y. A., Rennert, O. M., & Chrousos, G. P. (2007). Mitochondria as key components of the stress response. Trends in Endocrinology & Metabolism, 18(5), 190–198. 10.1016/j.tem.2007.04.004

McEwen, B. S. (1998). Protective and Damaging Effects of Stress Mediators. New England Journal of Medicine, 338(3), 171–179. 10.1056/NEJM199801153380307

McEwen, B. S. (2001). Plasticity of the hippocampus: Adaptation to chronic stress and allostatic load. Annals of the New York Academy of Sciences, 933, 265–277. 10.1111/j.1749-6632.2001.tb05830.x

McEwen, B. S., & Wingfield, J. C. (2003). The concept of allostasis in biology and biomedicine. Hormones and Behavior, 43(1), 2–15. 10.1016/S0018-506X(02)00024-7

Mehaisen, G. M. K., Eshak, M. G., Elkaiaty, A. M., Atta, A.-R. M. M., Mashaly, M. M., & Abass, A. O. (2017). Comprehensive growth performance, immune function, plasma biochemistry, gene expressions and cell death morphology responses to a daily corticosterone injection course in broiler chickens. PLoS ONE, 12(2), e0172684. 10.1371/journal.pone.0172684

Metcalfe, N. B., & Monaghan, P. (2013). Does reproduction cause oxidative stress? An open question. Trends in Ecology & Evolution, 28(6), 347–350. 10.1016/j.tree.2013.01.015

Monaghan, P. (2010). Telomeres and life histories: The long and the short of it. Annals of the New York Academy of Sciences, 1206(1), 130–142. 10.1111/j.1749-6632.2010.05705.x

Monaghan, P., & Haussmann, M. F. (2015). The positive and negative consequences of stressors during early life. Early Human Development, 91(11), 643–647. 10.1016/j.earlhumdev.2015.08.008

Murone, J., DeMarchi, J. A., & Venesky, M. D. (2016). Exposure to Corticosterone Affects Host Resistance, but Not Tolerance, to an Emerging Fungal Pathogen. PloS One, 11(9), e0163736. 10.1371/journal.pone.0163736

Names, G. R., Schultz, E. M., Krause, J. S., Hahn, T. P., Wingfield, J. C., Heal, M., Cornelius, J. M., Klasing, K. C., & Hunt, K. E. (2021). Stress in paradise: Effects of elevated corticosterone on immunity and avian malaria resilience in a Hawaiian passerine. The Journal of Experimental Biology, 224(20), jeb242951. 10.1242/jeb.242951

O’Connor, C. M., Gilmour, K. M., Arlinghaus, R., Van Der Kraak, G., & Cooke, S. J. (2009). Stress and Parental Care in a Wild Teleost Fish: Insights from Exogenous Supraphysiological Cortisol Implants. Physiological and Biochemical Zoology, 82(6), 709–719. 10.1086/605914

Oluwagbenga, E. M., Tetel, V., Tonissen, S., Karcher, D. M., & Fraley, G. S. (2023). Chronic treatment with glucocorticoids does not affect egg quality but increases cortisol deposition into egg albumen and elicits changes to the heterophil to lymphocyte ratio in a sex-dependent manner. Frontiers in Physiology, 14, 1132728. 10.3389/fphys.2023.1132728

Ouyang, J. Q., Muturi, M., Quetting, M., & Hau, M. (2013). Small increases in corticosterone before the breeding season increase parental investment but not fitness in a wild passerine bird. Hormones and Behavior, 63(5), 776–781. 10.1016/j.yhbeh.2013.03.002

Patterson, S. H., Hahn, T. P., Cornelius, J. M., & Breuner, C. W. (2014). Natural selection and glucocorticoid physiology. Journal of Evolutionary Biology, 27(2), 259–274. 10.1111/jeb.12286

Reichert, S., & Stier, A. (2017). Does oxidative stress shorten telomeres in vivo? A review. Biology Letters, 13(12), 20170463. 10.1098/rsbl.2017.0463

Remage-Healey, L., & Romero, L. M. (2001). Corticosterone and insulin interact to regulate glucose and triglyceride levels during stress in a bird. American Journal of Physiology-Regulatory, Integrative and Comparative Physiology, 281(3), R994–R1003. 10.1152/ajpregu.2001.281.3.R994

Ricklefs, R. E., & Wikelski, M. (2002). The physiology/life-history nexus. Trends in Ecology & Evolution, 17(10), 462–468. 10.1016/S0169-5347(02)02578-8

Robin, J.-P., Fayolle, C., Decrock, F., Thil, M.-A., Côté, S. D., Bernard, S., & Groscolas, R. (2001). Restoration of body mass in King Penguins after egg abandonment at a critical energy depletion stage: Early vs late breeders. Journal of Avian Biology, 32(4), 303–310. 10.1111/j.0908-8857.2001.320403.x

Romero, L. M., & Wikelski, M. (2001). Corticosterone levels predict survival probabilities of Galápagos marine iguanas during El Niño events. Proceedings of the National Academy of Sciences, 98(13), 7366–7370. 10.1073/pnas.131091498

Romero, L. M. (2004). Physiological stress in ecology: Lessons from biomedical research. Trends in Ecology & Evolution, 19(5), 249–255. 10.1016/j.tree.2004.03.008

Romero, L. M., Strochlic, D., & Wingfield, J. C. (2005). Corticosterone inhibits feather growth: Potential mechanism explaining seasonal down regulation of corticosterone during molt. Comparative Biochemistry and Physiology Part A: Molecular & Integrative Physiology, 142(1), 65–73. 10.1016/j.cbpa.2005.07.014

Romero, M. L., & Butler, L. K. (2007). Endocrinology of Stress. International Journal of Comparative Psychology, 20(2). 10.46867/ijcp.2007.20.02.15

Romero, L. M., Dickens, M. J., & Cyr, N. E. (2009). The reactive scope model—A new model integrating homeostasis, allostasis, and stress. Hormones and Behavior, 55(3), 375–389. 10.1016/j.yhbeh.2008.12.009

Sapolsky, R. M., Romero, L. M., & Munck, A. U. (2000). How do glucocorticoids influence stress responses? Integrating permissive, suppressive, stimulatory, and preparative actions. Endocrine Reviews, 21(1), 55–89. 10.1210/edrv.21.1.0389

Sapolsky, R. M. (2021). Glucocorticoids, the evolution of the stress-response, and the primate predicament. Neurobiology of Stress, 14, 100320. 10.1016/j.ynstr.2021.100320

Saraux, C., Viblanc, V. A., Hanuise, N., Maho, Y. L., & Bohec, C. L. (2011). Effects of Individual Pre-Fledging Traits and Environmental Conditions on Return Patterns in Juvenile King Penguins. PLOS ONE, 6(6), e20407. 10.1371/journal.pone.0020407

Schielzeth, H., Dingemanse, N. J., Nakagawa, S., Westneat, D. F., Allegue, H., Teplitsky, C., Réale, D., Dochtermann, N. A., Garamszegi, L. Z., & Araya-Ajoy, Y. G. (2020). Robustness of linear mixed-effects models to violations of distributional assumptions. Methods in Ecology and Evolution, 11(9), 1141–1152. 10.1111/2041-210X.13434

Schoenle, L. A., Zimmer, C., & Vitousek, M. N. (2018). Understanding Context Dependence in Glucocorticoid–Fitness Relationships: The Role of the Nature of the Challenge, the Intensity and Frequency of Stressors, and Life History. Integrative and Comparative Biology, 58(4), 777–789. 10.1093/icb/icy046

Schoenle, L. A., Zimmer, C., Miller, E. T., & Vitousek, M. N. (2021). Does variation in glucocorticoid concentrations predict fitness? A phylogenetic meta-analysis. General and Comparative Endocrinology, 300, 113611. 10.1016/j.ygcen.2020.113611

Sheriff, M. J., Dantzer, B., Delehanty, B., Palme, R., & Boonstra, R. (2011). Measuring stress in wildlife: Techniques for quantifying glucocorticoids. Oecologia, 166(4), 869–887. 10.1007/s00442-011-1943-y

Spée, M., Marchal, L., Thierry, A.-M., Chastel, O., Enstipp, M., Maho, Y. L., Beaulieu, M., & Raclot, T. (2011). Exogenous corticosterone mimics a late fasting stage in captive Adélie penguins (Pygoscelis adeliae). American Journal of Physiology-Regulatory, Integrative and Comparative Physiology, 300(5), R1241–R1249. 10.1152/ajpregu.00762.2010

Stearns, Stephen C, The Evolution Of Life Histories (Oxford, 1998; online edn, Oxford Academic, 31 Oct. 2023), 10.1093/oso/9780198577416.001.0001

Stier, A., Delestrade, A., Zahn, S., Arrivé, M., Criscuolo, F., & Massemin-Challet, S. (2014a). Elevation impacts the balance between growth and oxidative stress in coal tits. Oecologia, 175(3), 791–800. 10.1007/s00442-014-2946-2

Stier, A., Viblanc, V. A., Massemin-Challet, S., Handrich, Y., Zahn, S., Rojas, E. R., Saraux, C., Le Vaillant, M., Prud’homme, O., Grosbellet, E., Robin, J., Bize, P., & Criscuolo, F. (2014b). Starting with a handicap: Phenotypic differences between early- and late-born king penguin chicks and their survival correlates. Functional Ecology, 28(3), 601–611. 10.1111/1365-2435.12204

Stier, A., Schull, Q., Bize, P., Lefol, E., Haussmann, M., Roussel, D., Robin, J.-P., & Viblanc, V. A. (2019). Oxidative stress and mitochondrial responses to stress exposure suggest that king penguins are naturally equipped to resist stress. Scientific Reports, 9(1), 8545. 10.1038/s41598-019-44990-x

Taff, C. C., Zimmer, C., & Vitousek, M. N. (2018). Efficacy of negative feedback in the HPA axis predicts recovery from acute challenges. Biology Letters, 14(7), 20180131. 10.1098/rsbl.2018.0131

Torres-Medina F, Cabezas S, Marchant TA, Wikelski M, Romero LM, Hau M, Carrete M, Tella JL, Blas J. Corticosterone implants produce stress-hyporesponsive birds. Journal of experimental biology. 2018 Oct 11;221(Pt 19):jeb173864. doi: 10.1242/jeb.173864.

Viblanc, V. A., Bize, P., Criscuolo, F., Le Vaillant, M., Saraux, C., Pardonnet, S., Gineste, B., Kauffmann, M., Prud’homme, O., Handrich, Y., Massemin, S., Groscolas, R., & Robin, J.-P. (2012a). Body Girth as an Alternative to Body Mass for Establishing Condition Indexes in Field Studies: A Validation in the King Penguin. Physiological and Biochemical Zoology, 85(5), 533–542. 10.1086/667540

Viblanc, V. A., Smith, A. D., Gineste, B., & Groscolas, R. (2012b). Coping with continuous human disturbance in the wild: Insights from penguin heart rate response to various stressors. BMC Ecology, 12(1), 10. 10.1186/1472-6785-12-10

Viblanc, V. A., Gineste, B., Robin, J.-P., & Groscolas, R. (2016). Breeding status affects the hormonal and metabolic response to acute stress in a long-lived seabird, the king penguin. General and Comparative Endocrinology, 236, 139–145. 10.1016/j.ygcen.2016.07.021

Viblanc, V. A., Schull, Q., Cornioley, T., Stier, A., Ménard, J.-J., Groscolas, R., & Robin, J.-P. (2018). An integrative appraisal of the hormonal and metabolic changes induced by acute stress using king penguins as a model. General and Comparative Endocrinology, 269, 1–10. 10.1016/j.ygcen.2017.08.024

Vitousek, M. N., Taff, C. C., Ryan, T. A., & Zimmer, C. (2019). Stress Resilience and the Dynamic Regulation of Glucocorticoids. Integrative and Comparative Biology, 59(2), 251–263. 10.1093/icb/icz087

Voellmy, I. K., Goncalves, I. B., Barrette, M.-F., Monfort, S. L., & Manser, M. B. (2014). Mean fecal glucocorticoid metabolites are associated with vigilance, whereas immediate cortisol levels better reflect acute anti-predator responses in meerkats. Hormones and Behavior, 66(5), 759–765. 10.1016/j.yhbeh.2014.08.008.

Voituron, Y., Josserand, R., Le Galliard, J.-F., Haussy, C., Roussel, D., Romestaing, C., & Meylan, S. (2017). Chronic stress, energy transduction, and free-radical production in a reptile. Oecologia, 185(2), 195–203. 10.1007/s00442-017-3933-1

Warne, J. P. (2009). Shaping the stress response: Interplay of palatable food choices, glucocorticoids, insulin and abdominal obesity. Molecular and Cellular Endocrinology, 13th Conference on the Adrenal Cortex (Adrenal 2008), 300(1), 137–146. 10.1016/j.mce.2008.09.036

Weimerskirch, H., Stahl, J. C., & Jouventin, P. (1992). The breeding biology and population dynamics of King Penguins Aptenodytes patagonica on the Crozet Islands. Ibis, 134(2), 107–117. 10.1111/j.1474-919X.1992.tb08387.x

Wilbourn, R. V., Moatt, J. P., Froy, H., Walling, C. A., Nussey, D. H. and Boonekamp, J. J. (2018). The relationship between telomere length and mortality risk in non-model vertebrate systems: a meta-analysis. Philosophical transactions of the Royal Society of London. Series B, Biological sciences 373, 20160447–9. 10.1098/rstb.2016.0447

Wingfield, J. C., Donna L. Maney, Creagh W. Breuner, Jacobs, J. D., Sharon Lynn, Ramenofsky, M., & Ralph D. Richardson. (1998). Ecological Bases of Hormone-Behavior Interactions: The “Emergency Life History Stage.” American Zoologist, 38(1), 191–206. http://www.jstor.org/stable/4620131

Wingfield, J.C. and Romero, L.M. (2000), Adrenocortical Responses to Stress and Their Modulation in Free-Living Vertebrates. Comprehensive Physiology, 2000: 211–234. 10.1002/j.2040-4603.2000.tb01516.x

Wingfield, J. C., & Kitaysky, A. S. (2002). Endocrine Responses to Unpredictable Environmental Events: Stress or Anti-Stress Hormones?. Integrative and Comparative Biology, 42(3), 600–609. 10.1093/icb/42.3.600

Wingfield, J. C., & Sapolsky, R. M. (2003). Reproduction and Resistance to Stress: When and How. Journal of Neuroendocrinology, 15(8), 711–724. 10.1046/j.1365-2826.2003.01033.x

Zimmer, C., Taff, C. C., Ardia, D. R., Ryan, T. A., Winkler, D. W., & Vitousek, M. N. (2019). On again, off again: Acute stress response and negative feedback together predict resilience to experimental challenges. Functional Ecology, 33(4), 619–628. 10.1111/1365-2435.13281

Zimmer, C., Taff, C. C., Ardia, D. R., Rose, A. P., Aborn, D. A., Johnson, L. S., & Vitousek, M. N. (2020). Environmental unpredictability shapes glucocorticoid regulation across populations of tree swallows. Scientific Reports, 10, 13682. 10.1038/s41598-020-70161-4

